# Niche formation and metabolic interactions result in stable diversity in a spatially structured cyanobacterial community

**DOI:** 10.1101/2022.12.13.520286

**Authors:** Sarah J.N. Duxbury, Sebastien Raguideau, Kelsey Cremin, Jerko Rosko, Mary Coates, Kieran Randall, Jing Chen, Christopher Quince, Orkun S. Soyer

## Abstract

Understanding how microbial communities maintain stable compositional diversity is key for predicting community function. Studies from species pairwise interactions and synthetic communities indicate that metabolic interactions and spatial organisation can influence coexistence, but the relevance of these factors in more complex communities is unclear. Model systems often lack multi-species complexity, thereby making it difficult to study community diversity temporally. Here we used a spatially-organised cyanobacterial enrichment community to investigate compositional diversity and its stability. Over a year of passaging in media without significant carbon source, we found that the community maintains relatively high diversity, with 17 co-existing bacterial species. Using short and long read shotgun metagenomics sequencing from different time point samples, we have reconstructed complete genomes. Genomic annotation of these species revealed complementary metabolic functions involving carbon breakdown and vitamin biosynthesis suggesting interactions amongst community members. Using isolated species, we provide experimental support for carbon provision through cyanobacterial slime and growth on the component substrates by representative members of the Proteobacteria and Actinobacteriota phyla. Additionally, we experimentally show vitamin provision and uptake between prototrophic and auxotrophic members. We also found genomic capability for (an)oxygenic photosynthesis and sulfur cycling in several species. We show consistent formation of oxygen gradients across ‘photogranule’ structures, supporting niches that can sustain these specific metabolic functions. These findings indicate that spatial niche formation and metabolic interactions enable maintenance of community compositional stability and diversity.

**SIGNIFICANCE STATEMENT:** Microbes exist as species-diverse communities in nature and understanding their stability is an open challenge in microbial ecology. We established and maintained a spatially-organised, photosynthetic microbial community from a freshwater reservoir through long-term culturing in laboratory medium. We found that this community maintained a taxonomically-diverse set of 17 bacterial species. Combining genomic and physiological assays, we characterised a novel filamentous cyanobacterium capable of carbohydrate-rich ‘slime’ secretion supporting growth of other microbes. We predict inter-species vitamin exchanges and identify sulfur cycling and alternative types of photosynthesis that are likely to be favoured in oxygen-free zones identified within the spatial structures. Our findings indicate that metabolic interactions and spatial structures can enable stable microbial coexistence in natural ecosystems.

## INTRODUCTION

Understanding the assembly and stability of species composition in microbial communities is key for predicting community function. Experimental studies involving enrichment of environmental samples in defined culture media indicate that assembled community composition and diversity in terms of higher-level taxonomy, can be related to nutritional richness of the environment (1, 2). For the experimental assembly of relatively low diversity microbiomes, the extent and maintenance of microbial diversity is suggested to relate to metabolic cross-feeding and the production of secondary metabolites for growth in minimal medium supplemented with carbon sources (2-5). Full genomic annotations and experimental verification of species-specific metabolite exchanges, however, remain limited.

Another factor that can influence diversity and stability in a community is spatial organisation (6, 7). Certain natural spatially-organised microbial communities are indicated to maintain taxonomic stability over time, in both environmental and host-associated contexts (7-9). Metabolic gradients found within some of these communities (10-14) are hypothesised to be important for microbial interactions and community composition (12, 15). In a laboratory-based single species culture, spatial stratification of the environment can allowed emergence of different strains maintained through competitive trade-offs (negative frequency-dependent selection) (16).

Despite these indicative results, the emergence of spatial organisation and metabolic interactions in a microbial community and their contribution to determining species composition and stability remain to be systematically studied. Current model systems often lack spatial structure and multi-species complexity, contrasting with natural communities (9, 17, 18). Studies on spatial systems have focused mainly on biofilms composed of one or two species (19-21) or have studied species succession during biofilm assembly (17, 18). Thus, temporal stability of final community composition in presence of spatial organisation remains under-studied. Mechanistic studies of metabolic interactions usually involve “bottom-up” approaches (3, 22-24), using culturable microbial pairs or specific microbial groups to predict interactions and stability at the community level. It is unclear to what extent the outcomes of such pairwise interactions, in homogenous environments, will be predictive for spatially-organised natural communities.

In order to address the maintenance of microbial diversity, we developed here a spatially-organised, cyanobacterium-dominated enrichment microbial community and investigated temporal compositional stability and metabolic interactions within it. Over one year of serial passaging in media without any significant carbon source, we found that this spatially-organised community maintains relatively high diversity, with 17 co-existing species, some with strain diversity. Genomic annotation of these species revealed distinct metabolic functions of oxygenic and anoxygenic photosynthesis, and metabolic interactions involving carbon and vitamin sharing and, sulfur cycling. In line with these predictions, we found experimental support for carbon exchange between the dominant cyanobacterial species and other species through carbon substrate secretion, using isolated members of the community. We detected vitamin prototrophy and auxotrophy respectively for isolated species and growth promotion of an auxotroph through vitamin sharing. We also found formation of consistent anoxic microenvironments within the spatial structures formed by the community, providing support for predicted anoxygenic photosynthesis and anoxic sulfur cycling reactions. Taken together, these findings indicate that a spatially-organised, photosynthetic community can develop into a temporally stable microcosm with spatial niches supporting metabolic functions and interactions enabling stability and diversity.

## RESULTS

### A cyanobacterial enrichment culture displays reproducible spatial structure formation

We collected water samples from a local freshwater reservoir and maintained these in a minimal medium lacking a significant carbon source (see *Methods*). After an initial irregular passaging period, we established a regular, serial passaging regime with a 1 in 200 dilution approximately every 35 days (Fig. 1A). We found repeatable spatial structure formation in these cultures over passages, consisting of both spherical and more irregularly shaped granules (ranging from mm to cm scale), as well as aggregates and connected biofilms (Fig. 1B). We refer to these structures collectively as ‘cyanobacterial granules’. Microscopy analyses showed these structures to be dominated by a filamentous cyanobacterium, that displays a characteristic gliding motility (25) similar to that seen in other filamentous cyanobacteria (26).

**Figure 1.**
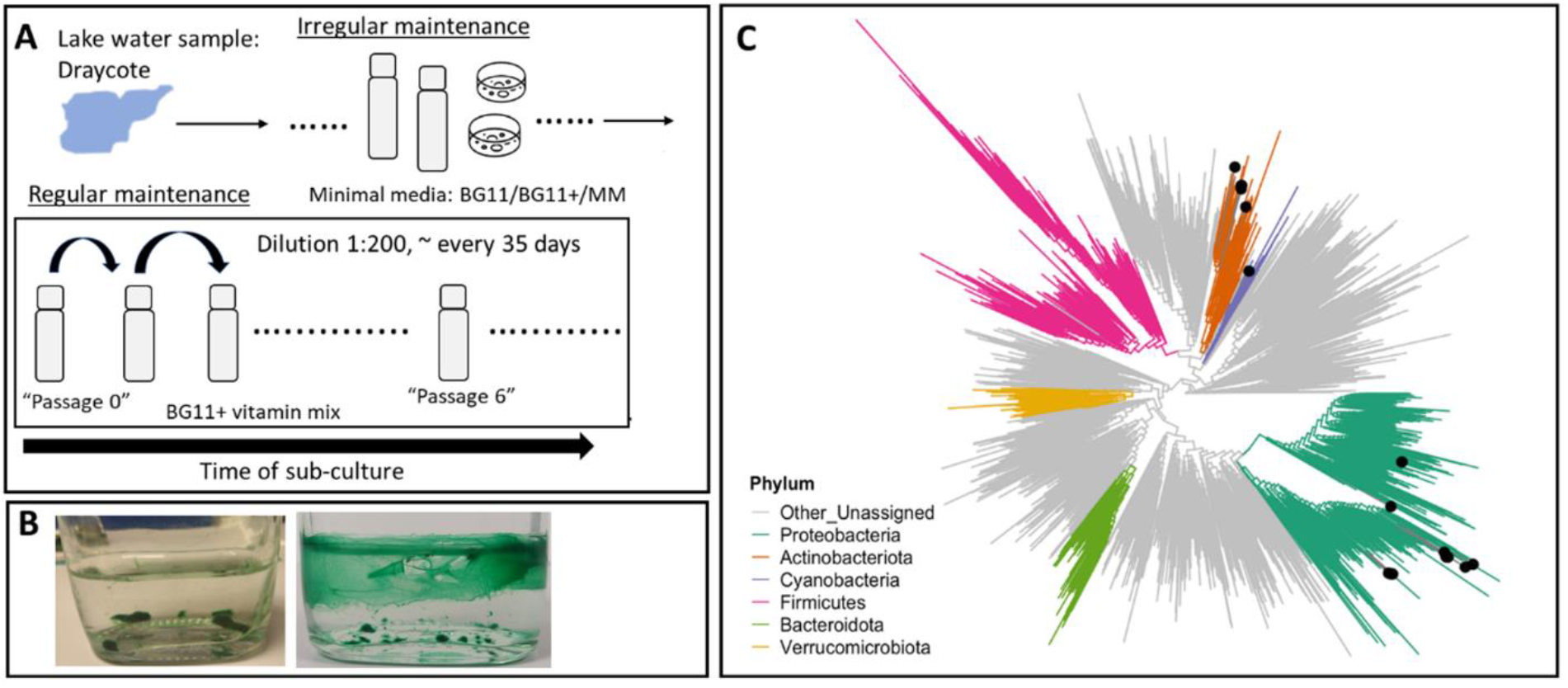
Culturing schematic, structure formation and taxonomic placement of granule-forming freshwater-derived phototrophic communities. **(A)** Schematic showing culturing regime from natural samples, followed by irregular culture maintenance in bottles and on agar plates prepared with minimal media. A regular laboratory serial passaging regime in culture bottles was initiated, for passages P0 to P6 that were sequenced for microbial community composition. See *Methods* for full details. **(B)** Representative images of the structured community. The image on the left show granules of various sizes and on the right, filamentous bundles and biofilms attaching to the base and walls of the glass bottle. **(C)** Phylogenetic tree showing taxonomic placement of the identified bacterial species in the community along with select phyla from the GTDB database (see *Methods*). The species from the community are highlighted with black dots, whilst colours indicate different phyla as listed in the legend. 16 species are presented on the tree, described in Table 1. For a maximum likelihood based phylogenetic tree of the cyanobacteria clade, including the novel cyanobacterium found in the system, see Fig S2.

### Cyanobacterial granules harbour a diverse, mid-complexity microbial community

From five passages ((P)assage P0 and P3-6) within a regular sub-culturing regime, we extracted DNA from mature cultures, performed short-read Illumina sequencing, and co-assembled raw sequencing data (see *Methods*). The sample from passage 1 was collected at an earlier culture age, therefore was analysed separately (Fig. S1), whilst data was not collected for Passage 2. Serial passaging of the community was extended (see *SI Appendix*) and a sample was collected from Passage 20 and analysed separately. We also characterised community composition of a sample prior to Passage 0 from the irregular culturing period (see *Methods* and Fig. S1).

**Table 1.**
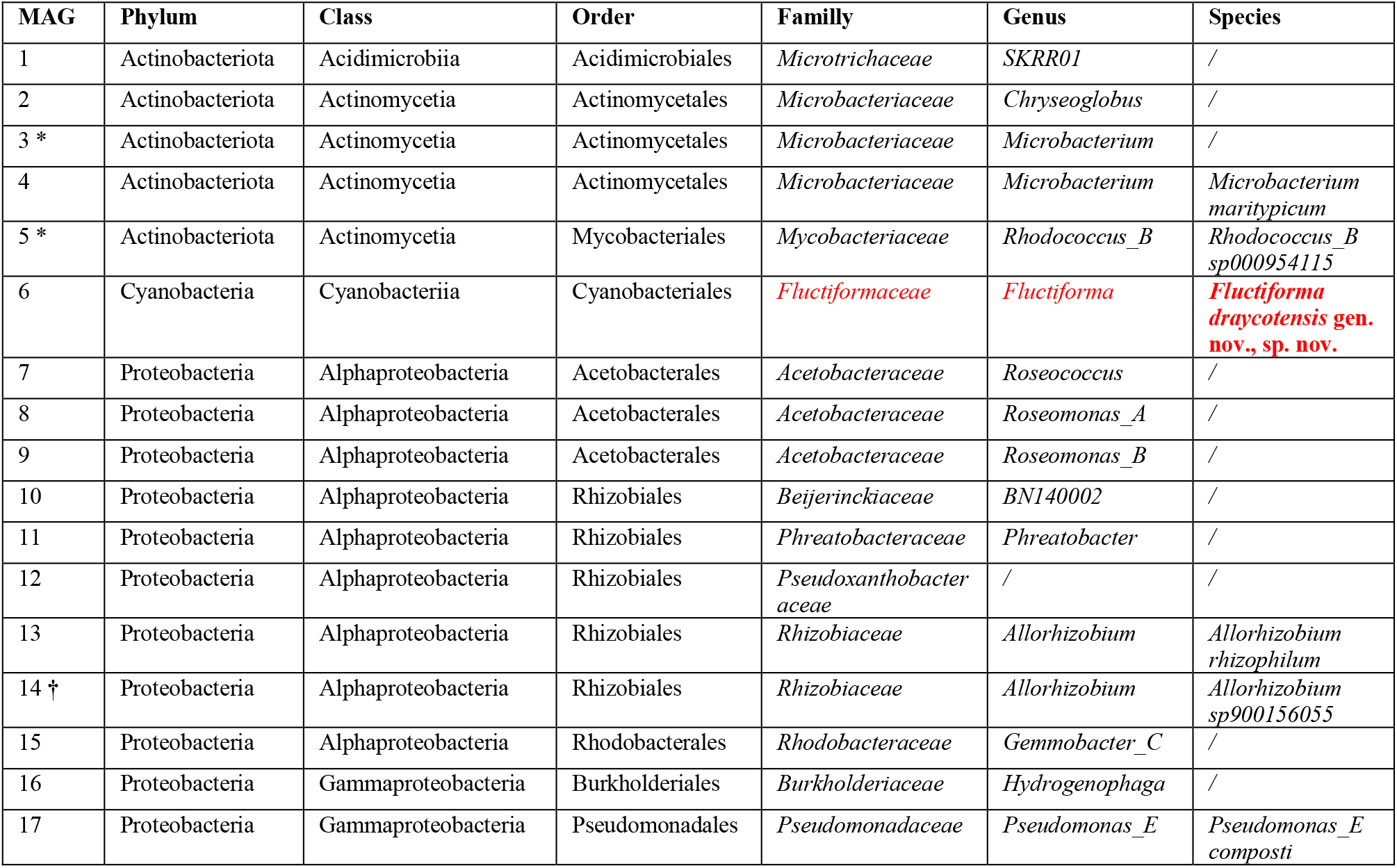
Taxonomy of bacterial species (MAGs) characterised in the structured cyanobacterial community. Assigned taxonomy of the final set of MAGs (metagenome assembled genomes) obtained from Illumina short-read and PacBio long-read shotgun metagenomics. MAGs only present in short-read sequencing are indicated by an ‘**\***’ in the ‘MAG’ column. A MAG that was only identified in long-read sequencing is indicated by a ‘†’. The highest taxonomic resolution is presented for each bin. The novel cyanobacterial species in our system is assigned by GTDB-Tk to an automatically generated family called ‘JAAUUE01’ (see main text for further discussion of the phylogeny of this species). We propose a novel taxonomic classification for this family and species, highlighted in red.

In all samples, we found the same set of 16 metagenome assembled genomes (MAGs) from short-read sequence data. For each sample, above 99% of raw sequence data mapped to the assembly, with sequence data mapped to MAGs varying between 82 – 99% across samples (see *Supplementary file 1*). This indicates that assembly and binning effort was highly successful at characterising the diversity of bacterial species extracted from this structured cyanobacterial community. We taxonomically assigned the MAGs using the genome taxonomy database (GTDB) phylogenetic placement toolkit (27) (see *Methods*, Fig. 1C and Table 1). Besides the mentioned cyanobacterium, the remaining 15 species were distributed in the phyla Actinobacteriota (5 species) and Proteobacteria (10 species) (Fig. 1C and Table 1). The latter group was spread between the alpha-(8 species) and gamma-Proteobacteria (2 species, *Pseudomonas E. composti* and a species of the *Hydrogenophaga* genus). The same species composition was also found in Passages 1 and 20 and in the early sample prior to Passage 0, with a greater enrichment of the cyanobacterium for younger culture age samples (Fig. S1). We have also noted that the system was generally robust to cryo-preservation (in 10% v/v glycerol), as cryo-revived community samples developed granules and showed the same species composition with comparable coverages to a similar aged community sample prior to cryopreservation (Table S1).

Most MAGs were taxonomically close to either a cultured species or to an uncultured MAG found on GTDB, with species-level novelty (see Table 1). The cyanobacterium had only one close homolog on GTDB, a single MAG obtained from sampling of extant stromatolites (28). Further analysis of the phylogeny of these two cyanobacteria (see *Methods*) showed that they form a novel family-level clade within the Cyanobacteriales order (Fig. S2). Based on its geographical origin and characteristic motility, we propose the name *Fluctiforma draycotensis* gen. nov., sp. nov. for the species found in our system and suggest a family name of *Fluctiformaceae*.

### Cyanobacterial granule community has stable species and strain composition

As a proxy for assessing stability of the community, we analysed coverage of each MAG across passages P0 and 3-6, normalised per Giga-base pair (Gbp) of sequencing to avoid biases from differences in sample sizes. This showed a qualitatively stable community composition over time, when analysed at the order level (Fig. 2A). This stable composition was similar to an early sample and the samples collected for Passages 1 and 20 (Fig S1). At the species level, some of the coverages showed correlations with passage number (Fig. 2B), but these were not statistically significant (p > 0.05; Pearson’s Correlation with Benjamini-Hochberg correction) (Table S2). *F. draycotensis* was found in much higher abundance than all other bins (average coverage 98-fold per Gbp across Passages 0 and 3-6). Using STRONG (29), a method for strain inference from longitudinal datasets, we identified two strains (‘haplotypes’) within each of the three species assigned to the *Allorhizobium* and *Chryseoglobus* genera, and the *Pseudoxanthobacteraceae* family (Table 1). All six strains were detected in all analysed passage samples, but for each species, one of the two strains was dominant (Fig. S3). Taken together, these results show a stable overall composition within the granule community over a one-year period, with relatively high taxonomic diversity and co-existence at both species and strain levels.

**Figure 2.**
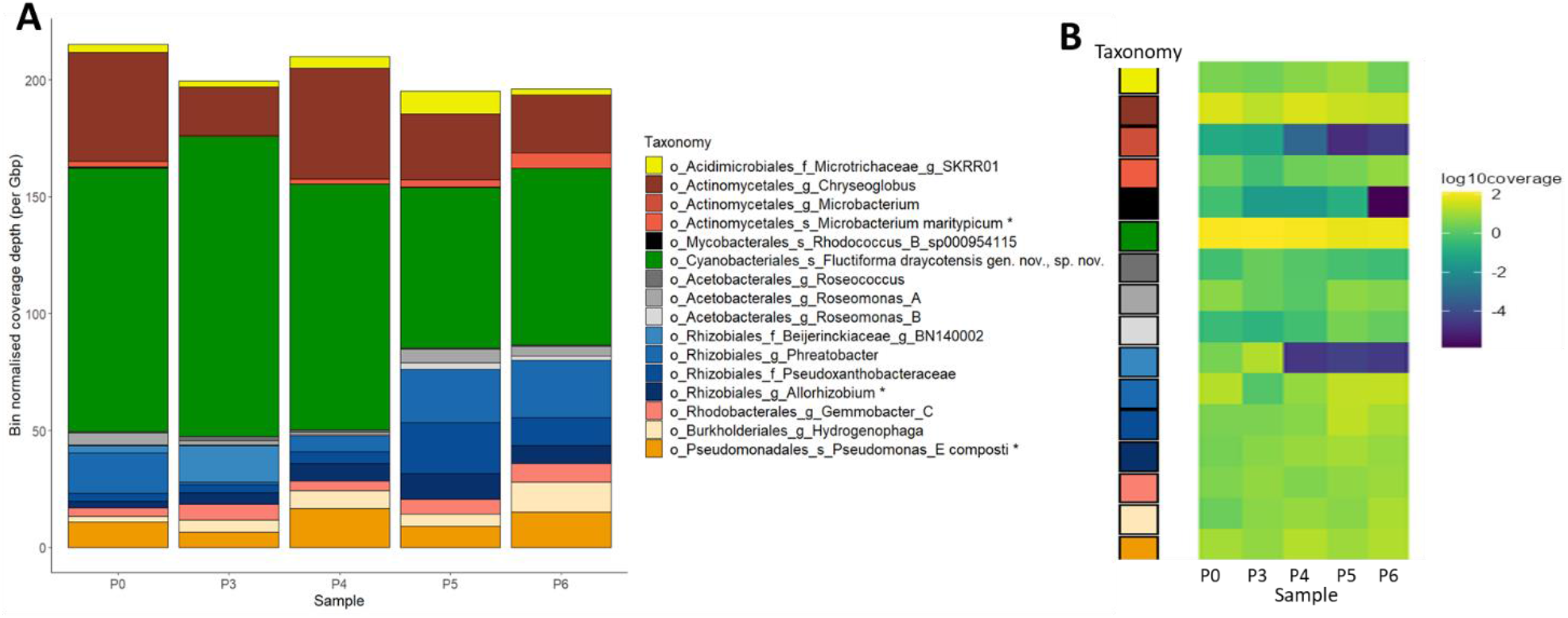
Coverage of taxonomic bins over serial passages. (**A**) Community composition, i.e. species coverage, over sub-culture passages (samples), colour-grouped at the taxonomic order level (listed in Table 1 and shown on the legend) for 16 species characterised from short-read Illumina sequencing. Colour shades indicate MAGs within orders and the highest taxonomic resolution is described in the legend on the figure. ‘o’ = order; ‘f’ = family; ‘g’ = genus; ‘s’ = species. Asterisks indicate species that were experimentally isolated. Passage number is labelled on the x-axis, denoting different samples. Data from Passage 1 is presented in Figure S1. The y-axis shows bin normalised coverage depth per Gbp of sequencing. (**B**) Same coverage data presented as in (A) but per individual bin and shown as log10-transformed. Taxonomy of bins is colour-coded as in (A). For Passage 6, no coverage of the *Rhodococcus_B_sp000954115* bin was detected so a pseudo-coverage value of 1.37e-06 (one order of magnitude lower than the smallest recorded coverage value) was applied to that Passage 6 bin (see Table S2).

### Long-read sequencing provides circular genomes of community members

To improve characterisation of community members, we used high molecular weight DNA extractions and long-read PacBio Hifi sequencing of two community samples (P0 and P7) (see *Methods*). Co-assembly of sequence data identified 14 MAGs in common with the short-read approach (Table 1 and S3), including resolution of an additional *Allorhizobium* species and two MAGs of the *Allorhizobium rhizophilum* species corresponding to different strains. This latter result is consistent with strain diversity predicted from short reads. Two species from the short-read sequencing (an unresolved species of the *Microbacterium* genus and *Rhodococcus B sp000954115*) were missing in the long-read sequencing, possibly due to low coverages (Fig. 2B).

For two species (*F. draycotensis* and the species from the *Chryseoglobus* genus), two separate complex (high variability) circular components were observed in the assembly graph. These appeared to be artifacts of high coverage as well as numerous wide overlapping regions between contigs. Strain diversity was explored as an explanation but no divergence in SCGs and 16S rRNA sequences nor strain diversity (via STRONG pipeline) were detected. Genomes were extracted through use of a consensus path algorithm resulting in the presented final genome sequences (see *Methods*). Eleven genomes showed 100% completeness based on presence of a set of single copy genes (SCGs) (see *Methods*), while the remaining had completeness at 75% or above (Table S3). Bin genome sizes ranged between 2.9Mb for the species from the *Chryseoglobus* genus to 6.8Mb for the species from the *Roseomonas A* genus, while GC content ranged between 44.3% for *F. draycotensis* to 71.3% for the species from the *Roseomonas B* genus. Twelve species were Gram-negative, including the cyanobacterium, and three were Gram-positive.

### Carbon fixation and slime-based carbon provision in the cyanobacterial community are predicted by genomic analyses and supported by experimental assays

The stable species coverage in this structured cyanobacterial community, together with the fact that our culture media lacked any significant carbon sources, implies the presence of carbon sharing among community members. To explore this hypothesis, we used the MAGs from both short- and long-read sequencing and analysed metabolic capabilities of the final set of 17 species, consisting of 14 species with both short and long-read sequences (combining *A. rhizophilum* strain genomes from long read data), one additional species found in long-read data (*A. sp900156055*) and two species with short-read sequences only (species from the *Microbacterium* genus and *Rhodococcus_B sp000954115*) (see *Methods*). This revealed majority presence of genes for central carbohydrate metabolism across all genomes (Fig. S4). As expected, the cyanobacterium *F. draycotensis* has the genes for the reductive pentose phosphate cycle (Calvin cycle) (Fig. S4) and the Rubisco genes *rbcL* and *rbcS*. Interestingly, the species from the *Pseudoxanthobacteraceae* family and *Hydrogenophaga* genus, also have both of these gene sets, indicating capability for carbon fixation in these two species. Three alternative autotrophic carbon fixation pathways were assessed but were not found to be complete in any of the genomes (Fig. S4).

These results show that only a few species in the system are capable of carbon fixation, suggesting that carbon provision by some of these species supports growth of the other species in the community. In particular, *F. draycotensis* gliding motility is associated with secreted ‘slime’ (exopolysaccharide), which is shown to be essential for gliding motility (26) and can have a diverse composition of monosaccharides including galacturonate and galactose (30). We found that galacturonate, glucuronate, and galactonate degradation pathways were above 75% complete in four species belonging to the alpha-Proteobacteria phylum (*Rhizobiales* and *Rhodobacterales* orders) and galactose degradation was present in three species of the Actinobacteria phylum (Fig. 3A). In support of bacterial slime degradation, we have also consistently observed bacterial attachment and growth on cyanobacterial slime (Fig S5).

**Figure 3.**
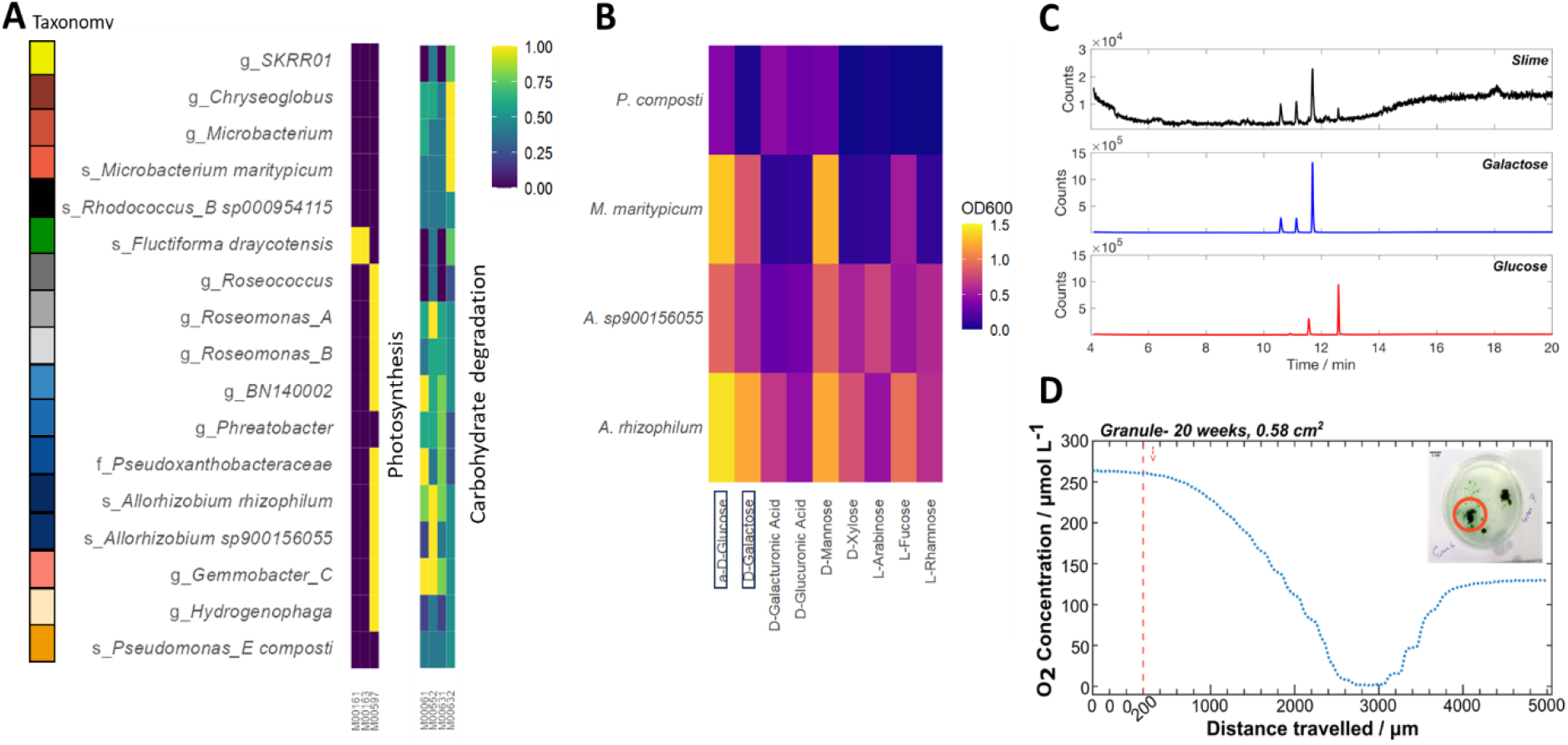
Functional potential and experimental verification for carbohydrate and energy metabolism and niche construction. **(A)** Completeness of select metabolic modules defined by KEGG orthology (KO) (see *Methods*). Bins are colour-coded by order (as in Fig. 2) and named by the highest taxonomic resolution resolved per bin (family, genus or species). Full taxonomy is given in Table 1. Module names are as follows. Photosynthesis: M00161 = PSII, M00163 = PSI, M00597 = anoxygenic PSII. Carbohydrate degradation: M00061 = glucuronate, M00552 = galactonate, M00631 = galacturonate, M00632 = galactose. **(B)** Growth of four isolated species on carbon substrates associated with cyanobacterial ‘slime’. Endpoint Optical Density (OD600) following 48 hours of growth is presented across carbon sources assayed in a Biolog PM1 microplate (see *Methods*). Those selected in this figure are based upon prior described components of cyanobacterial slime (30, 31). **(C)** GC chromatograms from GC-MS analysis of cyanobacterial ‘slime’ (exopolysaccharide) (top trace in black), compared with chromatograms of galactose and glucose standards (blue and red, respectively). The x-axis shows retention time. See Fig S6 for mass spectra. **(D)** Representative oxygen profile of a cyanobacterial granule, where 0 μm represents the starting position of the microelectrode probe at 1 mm above the granule, followed by the distance travelled with a timed z-axis step program (200 μm step, at each 20 sec point, unless otherwise stated). The red line shows the first descending step of the experiment. The inset image shows the granule prior to the oxygen profile measurement.

To further confirm these findings on predicted slime-based carbon exchange, we first extracted the ‘slime’ exopolysaccharide component from a community culture sample and used GC-MS chemical analyses to identify its monosaccharide components (see *Methods*). Comparing the GC chromatograms of the slime extract with profiles for a characterised set of monosaccharides, we identified galactose and glucose in the slime extract (Fig. 3C). While this analysis was based on peak retention profiles, we further confirmed the identity of these compounds using mass spectra (Fig S6).

We then attempted isolation of individual species from the cyanobacterial community to test growth on these and other carbon sources. Via dilution plating (see *Methods)*, we isolated *P. composti, A. rhizophilum, A. sp900156055* and *M. maritypicum*. Biolog growth assays were performed on a range of carbon sources used to support growth (see *Methods*), including the sugars for which degradation pathways were identified. We observed growth on galacturonate and glucuronate for *A. rhizophilum, A. sp900156055* and *P. composti* (Fig. 3B). Growth on galactose was seen for *A. rhizophilum, A. sp900156055* and *M. maritypicum* despite lack of complete biosynthesis pathways in their assembled genomes. In addition, growth, particularly for *A. rhizophilum, A. sp900156055* and *M. maritypicum*, was seen on several other sugars (glucose, xylose, mannose, fucose, arabinose) that are also described as components of cyanobacterial slime (30-32).

Additionally, we explored pathways for amino acid and nitrogen metabolism. Amino acids are not supplied directly in the growth medium, so biosynthesis is required. We did not find a stark variation in amino acid biosynthetic capacity across the genomes suggestive of complementary metabolic interactions (Fig S7). Regarding nitrogen metabolism, we did not find any genes for nitrogen fixation across the genomes, indicating that nitrate supply is essential in the medium.

### Vitamin biosynthesis and inter-species provision is predicted from genomic data and supported by isolate and community-level growth assays

Previous studies have shown that vitamin-based interactions occur between algae or cyanobacteria and other bacteria (33-35). Analysing the completeness of vitamin biosynthesis pathways in the presented genomes (Fig. S8), we identified two key vitamins – vitamin B7 (biotin) and B12 (cobalamin) – for which pathways were only fully complete in a single species: *P. composti* (see *SI Appendix* and Figures 4A and S9). The species *A. rhizophilum* has near-complete modules for all sections of the vitamin B12 pathway (Fig. S9A). All other species have partially complete B12 pathways, some also missing synthesis genes for key intermediates but with scavenging pathways complete, including *F. draycotensis* (Fig. S9A). For vitamin B7, *P. composti* showed complete presence of all biosynthetic genes, whereas several genes, particularly for the lower part of the synthesis pathway, were missing across all other species (Fig. 4A). To test for these contrasting predictions in vitamin prototrophy in isolated species from the community, we grew *P. composti* and *A. rhizophilum* in BG11+ medium supplemented with 0.8% w/v glucose, in the presence or absence of vitamin mix including vitamin B7. We found that *P. composti* could grow to high density with or without vitamin mix addition with average growth even slightly greater (1.07-fold) in the absence of vitamin mix (two-sample t-test: t_(4)_ = -3.33, p = 0.029) (Fig. 4B). Growth of *A. rhizophilum* was greatly limited in the absence of vitamin mix, with endpoint growth density reduced by 9.6-fold (one-way ANOVA: F_(3, 12)_ = 2581, p < 2e-16; Tukey pairwise p-adj = 2.7e-14) (Fig 4C). Supplementation of BG11+ medium containing 0.8% w/v glucose, with vitamin B7 or with supernatant collected from a grown culture of *P. composti* (see *SI Appendix)* restored growth of *A. rhizophilum* above that of the no vitamin mix treatment (Tukey pairwise p-adj = 2.7e-14; 3.7e-14 respectively) but below the level of full vitamin supplementation (p-adj = 4.2e-07; 1.4e-08 respectively). This shows that *A. rhizophilum* is auxotrophic for vitamin B7 and provision from *P. composti* biosynthesis can restore growth.

**Figure 4.**
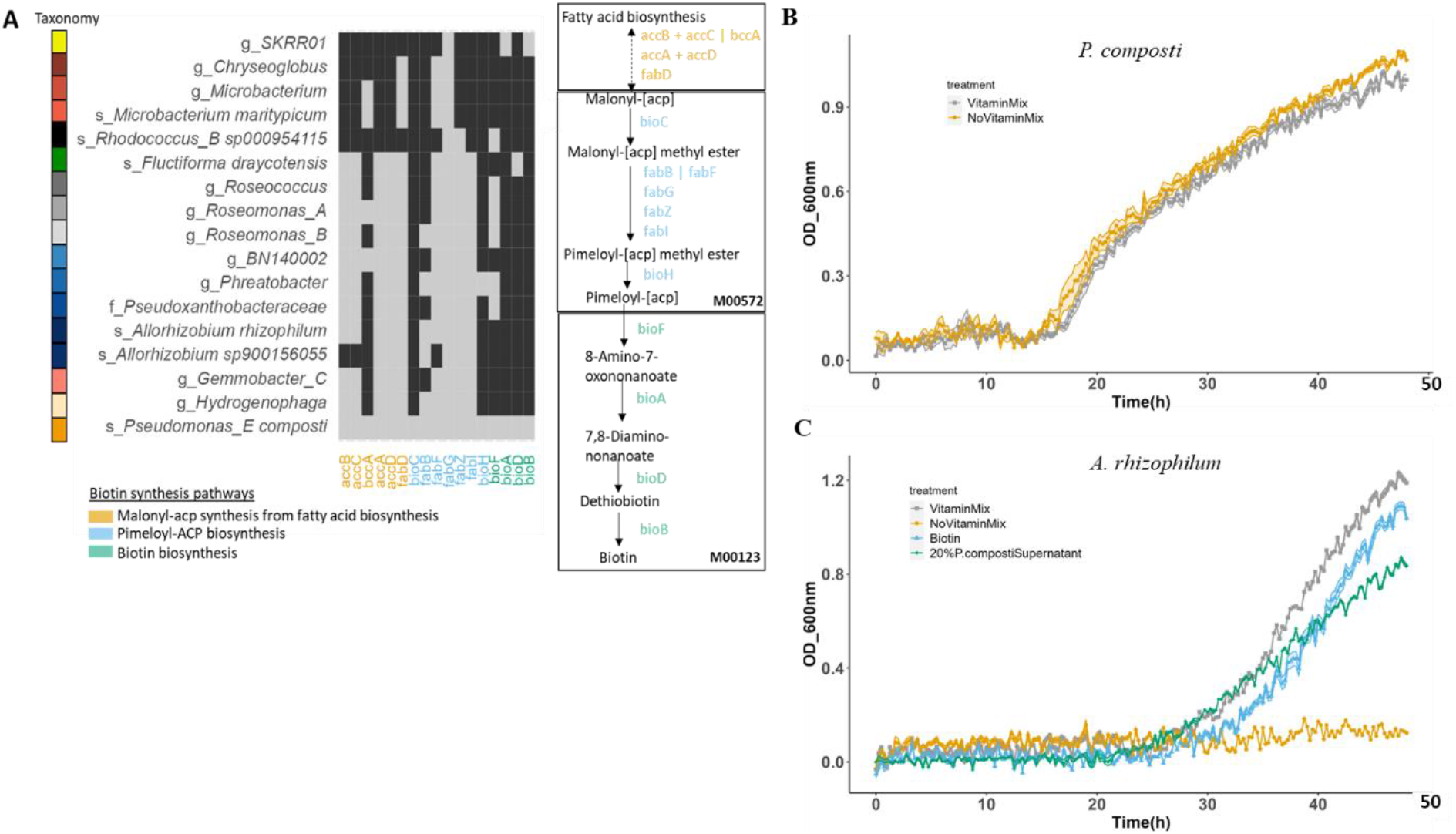
Functional analysis and experimental verification of vitamin B7 (biotin) biosynthesis capabilities and suggested inter-species provision. **(A)** Representation of completeness of biotin pathways defined by KOs (KEGG Orthologs). Heatmap plot shows presence (light grey) and absence (dark grey) of select genes, colour-coded by pathway, across the full set of 17 species. See Table 1 for full species names and taxonomy. See *Methods* for full details of gene annotation and analysis and *Supplementary file* 3 for KO analyses. Defined pathway steps and genes required for full synthesis of biotin are illustrated to match the x-axis of the heatmap, listing pathway genes present across at least one genome. On the pathway schematic, vertical bars separate alternative genes and plus symbols show required gene combinations. **(B)** Growth of *P. composti* in BG11+ vitamin mix medium supplemented with 0.8% w/v glucose, in the absence and presence of vitamin mix. N = 3 per treatment. **(C)** Growth of *A. rhizophilum* in BG11+ medium supplemented with 0.8% w/v glucose, in the absence and presence of vitamin mix or with separate supplementation of biotin, or *P. composti* supernatant mixed with BG11+ medium in a 20:80 volumetric ratio (1x BG11+ medium final concentration). N = 4 per treatment. Treatments are represented by different colours and symbols. Points plotted represent averages across replicates and shaded regions represent standard errors.

Overall, the patchiness of biosynthetic genes across the different species in our system suggests possible metabolic interactions mediated by vitamin B7 and B12, not only between the cyanobacterium and the other bacterial species, but among the latter as well. Observation that the community can re-grow and form granules in BG11+ medium without vitamin mix supplementation and over three serial growth passages (see *SI Methods* and Fig. S10), provides further evidence for exchange of essential vitamins among community members. Biomass however significantly varied between the two treatments (BG11+ medium with or without vitamin mix) when taking into account variation across passages (χ^2^_1_ = 8.357, p = 0.00384), with an average 1.6-fold reduction in biomass without vitamin mix addition.

### Oxygen gradients in the structured community indicate spatial metabolic niches

Genomic annotations confirmed the photosynthetic capability of *F. draycotensis*, as expected, but also revealed anoxygenic photosynthesis capability in nine species (Fig. 3A). Anoxygenic photosynthesis is an alternative type of photosynthesis whereby alternative electron donors such as hydrogen sulfide - are used, instead of water, resulting in no oxygen production (36).

In addition, several genomes contained genes involved in assimilation and cycling of sulfur (Fig. S11A). Two species, those from the *Hydrogenophaga* and BN140002, show capability of thiosulfate oxidation, whilst five species showed capability of sulfite oxidation (all species had a complete SOE complex). Several species, particularly *Pseudomonas_E. composti* and the species from the *Pseudoxanthobacteraceae* family showed capability for sulfate and sulfite reduction via the assimilatory pathway. In contrast, we did not find presence of dissimilatory sulfate reduction genes used in anaerobic respiration (37) across any of the genomes (Fig. S11A and C). Additionally, we found that *P. composti*, the species from the *Pseudoxanthobacteraceae* family and *F. draycotensis*, contain alkanesulfonate, thiosulphate and sulfate transport genes and genes for subsequent conversion to sulfite and polysulfide (Fig. S11A), These findings suggest interspecies sulfur exchanges via secretion and uptake, as well as the presence of a sulfur cycle in the system, through sulfate reduction to sulfite and hydrogen sulfide, and subsequent oxidation of these compounds – and related thiosulfate and sulfur (38, 39) – back to sulfate via the SOX and SOE complexes (Fig. S11B).

We hypothesised that co-existence of oxygenic and anoxygenic phototrophs as well as presence of sulfur oxidation and reduction was enabled due to formation of oxygen gradients. To test this, we used micron-scale electrochemical probes to measure oxygen across cyanobacterial granules. As expected from oxygenic photosynthesis of the cyanobacterial species (*F. draycotensis*), high oxygen levels were detected at the granule-exterior boundary (Fig. 3D and S12). Entering the granule, a steep inwardly decreasing oxygen gradient was detected with the granule core being anoxic. This oxygen gradient was measured consistently across a range of differently sized and aged granules (Fig. S12). Taken together, these findings suggest that distinct metabolic functions are located within oxic/anoxic microenvironments developing in cyanobacterial granules, thereby contributing to diversity and stability of community composition.

## DISCUSSION

We have shown that a spatially-structured cyanobacteria-dominated microbial community, kept in the laboratory, displays a mid-complexity, temporally stable species composition, significant taxonomic diversity, and stable species and strain co-existence. Our combined genomic analyses and experimental measurements indicate that species co-existence is driven by metabolic exchanges facilitating growth in minimal medium, and formation of spatial microenvironmental niches within the granules. Using short and long-read shotgun metagenomic sequencing enabled full genome assembly and annotation of metabolic functions, aspects that are missed from studies characterising microbial diversity at the level of the 16S rRNA gene (2, 3, 40) or from theoretical studies of community interactions (4, 5). Combined with species isolation and experimental verification of interactions, this system provides an example of linking microbial diversity with functional traits, an area still largely under-explored for nature-derived communities (41-43).

Overall, we note that the composition of the presented community has similarities, at higher taxonomic levels, to that found in previous studies of cyanobacterial aggregates and blooms (17, 44-48). In particular, alpha and gamma-Proteobacteria are highly abundant in blooms, and members of the orders Rhodobacterales, Rhizobiales, Burkholderiales and Pseudomonadales (49, 50), and the *Hydrogeonophaga* genus (44, 49) are commonly detected. Some phyla such as the Verrucomicrobiota and Planctomycetia that are implied to be enriched in blooms are missing in our system. It must be noted, however, that all previous studies to date were based on natural samples collected at a single or few time points across a bloom, while the presented analysis focussed on a nature-derived system kept in the laboratory over many passages. Thus, it is possible that there is a core cyanobacteria-associated community (i.e. a *cyanosphere*) that can be stably recovered under laboratory conditions, while there are also peripheral community members that may be lost over time due to shifted environmental conditions. In support, a core microbiome has recently been described to be recruited *in vitro* alongside model cyanobacterial species, derived from diverse bacterial consortia cultured from freshwater environments (51). Similar to our findings, bacteria from the phyla Bacteroidetes and Proteobacteria, specifically the orders Rhizobiales and Rhodobacterales, were commonly enriched.

We have identified both functional potential and experimental support for metabolic complementarities in the presented community, in terms of sulfur cycling, carbon and vitamin exchanges. These metabolic interactions are likely to provide a basis for symbioses, as previously implicated in aquatic environments and microbial mats (44, 52-54). Combined with the formation of micro-environmental niches, we demonstrate both species and strain coexistence through passages. Given that we observe long-term survival and biofilm growth of community cultures for several months in the absence of medium replenishment, we predict that metabolic exchanges are essential for community function. This community may therefore provide an example of a self-sustaining ecosystem (55).

Through experimental characterisation of cyanobacterial slime and growth of a set of isolated species on monosaccharides constituting the slime, we show that a key exchange is carbon-mediated. Excretion of carbon-containing metabolites from phototrophs has been suggested to influence heterotroph community assembly (17, 40, 56). For slime-producing cyanobacteria, such influence can arise through selectivity of bacteria that can attach to or degrade specific types of slime. Such selective mechanisms based on mucus is described in coral (57) and squid (58) species and suggested in the vertebrate gut (59).

Auxotrophies are common in microbial ecosystems, particularly aquatic ones, and are predicted to lead to species interactions that influence microbial community composition (54, 60, 61). Our data provides support for multiple potential interactions mediated by vitamins, based on patchiness of biosynthetic genes across different species in our enrichment photosynthetic community. Vitamin auxotrophies are especially abundant, such as for cobalamin, biotin and thiamine in algal species in various aquatic environments (62). Many bacteria across diverse environments also use or produce cobamides (60). As a result, extensive nutrient exchanges are predicted in multi-species communities. Combining with species isolation, we could provide experimental support for complementary auxotrophic and prototrophic functions for biotin synthesis predicted from genomic annotations in our community, whereby one species (*P. composti*) is predicted to synthesise biotin for the other community species. For vitamin B12 (cobalamin) synthesis, we predict from genomic data that two heterotrophs in the presented community are likely to be prototrophic. These findings align with functional predictions of vitamin B12 biosynthesis by epibionts and sharing with *Microcystis* species in bloom communities, supported by community re-growth in the absence of vitamin B12 (44), similar to re-growth of our community in the absence of vitamins. Additionally, core species associated with freshwater cyanobacteria *in vitro* have shown enrichment of vitamin B12 biosynthetic genes (51).

The presented system displays granule formation, therefore showing spatial organisation that may have contributed to the maintenance of diversity via the formation of microenvironmental niches (6). We have consistently quantified oxygen gradients in granules. These gradients support the genomically identified functional traits, indicating coexistence of oxygenic and anoxygenic phototrophs in the presented community, as well as sulfur cycling. Sulfur cycling has been predicted to occur in bloom communities associated with *Microcystis* species from functional annotation of assimilatory sulfur reduction and oxidation genes (44). Anoxygenic photosystems have been characterised in core bacterial species found to associate with model cyanobacterial species, suggesting a role for light energy capture in more energy-intensive metabolic pathways (51). Anoxygenic photosynthesis was first discovered in anaerobic bacteria (63), however it is also maintained in the presence of oxygen (in aerobic anoxygenic phototrophs (64)). It is known that this type of photosynthesis specialises on longer (above 700 nm) wavelengths (65), so the shading effects and oxygen gradients arising from structure formation could allow these species to scavenge higher wavelengths of light than those used by *F. draycotensis*. The oxygen-poor granule core is likely to harbour bacteria that perform the sulfur reduction reactions, generating sulfide that could be used by anoxygenic phototrophs as an energy source. Sulfide could also act as a substrate for the sulfur oxidation reactions. We were not able to consistently detect presence of a hydrogen sulfide gradient through the granules, however this could have been due to variable dynamics from diffusion processes within the granule or through balanced sulfide production and consumption. Future studies can be designed to confirm predictions for spatial organisation of microbes, using taxa-specific labelling approaches such as FISH (Fluorescence *In Situ* Hybridisation) (66).

The presented system provides an ideal model to test hypotheses on niche formation and species diversity, an important and open current area of investigation in microbial ecology (6, 67). In addition to combination of metagenomics and experimental assays, we envisage that development of meta-transcriptomics (48), proteomics (68) and metabolomics (17) tools will aid discovery of live community members and metabolic activity of the community. Overall, the medium complexity and cultivability of this presented microbial community provides a model for further studies on relations between spatial organisation and community function, stability and evolution.

## MATERIALS AND METHODS

For a detailed description of all experimental methods, see SI Appendix.

### Sample collection and culture maintenance

Freshwater samples were collected from Draycote Water Reservoir, Warwickshire, UK. Cultures were maintained in BG11+ medium (DSMZ medium reference number 1593), with addition of a vitamin mix (Table S6) but without carbon source addition. Cultures were grown under continuous 12h/12h light/dark cycles with white, fluorescent illumination of 14 – 20 µmol photons m^-2^ s^-1^ at room temperature under static conditions. For each sub-culture, we performed a 1 in 200 dilution by transferring 150 µl of re-suspended filamentous culture into a final volume of 30 ml of BG11+ vitamin mix (Fig. 1A).

### DNA extraction and sequencing of community samples

For shotgun sequencing, DNA was extracted using the Qiagen PowerSoil Pro kit (Hilden, Germany, Cat. No. 47014). DNA was sent to Novogene (Cambridge, UK) for Illumina NovaSeq paired-end sequencing. For PacBio HiFi sequencing, DNA extractions were performed by the Natural Environment Research Council (NERC) Environmental Omics Facility (NEOF) using the Macherey-Nagel NuceloBond HMW DNA kit. PacBio DNA libraries and sequencing were completed at the Centre for Genomics Research (CGR). Sequencing was performed on the Sequel II SMRT Cell in CCS run mode.

### Sequence assembly, binning and coverage analysis

Samples from Passage (P)0 and 3-6 were processed using the STRONG pipeline (29). Firstly, samples were co-assembled through metaSPAdes, resulting in a normal assembly graph and a high resolution assembly graph retaining strain diversity as a path in the graph. Regular assembly was then binned into MAGs. Per MAG, single copy core genes (SCGs) were used to extract the part of the high resolution assembly graph centered around these SCGs. Number of strains as well as path in the subgraphs were estimated by reconstructing the coverage in the graph for each sample of the time series. Strain specific SCGs as well as MAG and strain coverages were generated. Normalisation was calculated by sample sequencing depth (per Gbp). Raw coverage values per sample are presented in *Supplementary file 2*. Coverage of the 16 MAGs in samples from P1 and 20 were calculated as a second step posterior to the STRONG run. To do so, reads were mapped using bwa-mem (69), and coverage was obtained using samtools (1.17) (70) and bedtools (v2.25.0) (71). The STRONG pipeline is available online at: https://github.com/chrisquince/STRONG. Percentages of reads recruited by either assembly or MAGs were calculated by mapping reads to the assembly and for the latter, summing over contigs of the same MAGs. The PacBio HiFi samples were assembled using hifiasm-meta (72) and the resulting unitig assembly graphs were used in the downstream analyses. We taxonomically classified MAGs with GTDB-Tk v2.1.0 (73) and data version r207, using standard settings on GTDB-Tk (27).

### Genome annotations

Genome annotations were performed using the DFAST annotation platform (releases 1.2.15/1.2.18; (74)). Resulting protein sequences were annotated for KEGG orthologs (KOs) using KofamKOALA (release 102.0/103.0; (75)). Resulting KO lists from short and long-read data were concatenated to create a unique KO list (where possible) for each of the MAGs/bins identified from the PacBio data. MetQy package (version 1.1.0) (76) was used to analyse the completeness of KEGG modules as defined by KEGG orthology (KO) blocks in the KEGG database (77).

### Species isolation and physiological assays

Isolation of species was achieved using three types of agar with carbon source supplementation: BG11+ vitamin mix with 0.1% w/v glucose, yeast mannitol and BG11+ vitamin mix with 0.05% w/v riboflavin. Cryostocks were prepared with a final concentration of 15% v/v glycerol for the cultures grown in BG11+ media and with 30% v/v glycerol (as of (78)) for the culture grown in yeast mannitol medium. Cryostocks were stored at -80°C. DNA was extracted from cell pellets using the Qiagen PowerBiofilm kit (Cat. No. 24000-50) and stored at -80°C. Isolates were characterised via Sanger sequencing of the bacterial 16S rRNA V3-V4 gene region. The Biolog ‘PM1 96 Carbon Utilisation Assay’ (Biolog, Hayward, CA, USA) was used with growth measured via absorbance (optical density at 600 nm wavelength) at 30°C for 48 hours in a plate reader (CLARIOstar, BMG LABTECH GmbH, Ortenberg, Germany). Isolated *P. composti* and *A. rhizophilum* species were growth-profiled in BG11+ medium in the presence and absence of vitamin mix over replicates in the plate reader. *A. rhizophilum* was also profiled in BG11+ medium in the presence of only biotin supplementation (same concentration as in vitamin mix) or with supplementation of *P. composti* supernatant from a culture grown in BG11+ medium without vitamin mix. Supernatant and BG11+ medium were mixed in a 20:80 ratio.

### Slime extraction for GC-MS characterisation

Slime extraction was performed adapting methods previously described (30, 32). The final protocol used is available via protocols.io (79). A culture of the community equivalent to Passage 16 of the regular culturing regime was selected for extraction. A culture of 1.2 L was grown for 3 months for sufficient slime production, producing approximately 50 mg of lyophilised slime.

### Micron-scale probe measurement of oxygen gradients

Oxygen was measured using a Unisense OX-NP-710704 probe. The microsensor was mounted onto a Scientifica IVM-Single Axis manipulator to control the probe movement. Cyanobacterial granules were extracted from cyanobacterial community samples (see *SI Appendix* for culture details) and placed into a dish with a layer of BG11+ agar and filled with fresh BG11+ media to cover the granule and the tip. The agar bed served as a method to protect the end of the tip from damage when it passed through the granule, it also helped to anchor the granule in a set position. The probe tip was then positioned at the approximate cyanobacterial granule surface and retracted back 1 mm for the start of the experiment (this position corresponds to 0 μm on the x-axis in Fig. 3D). Following an initial 60-second rest period for stabilising the signal, the probe was then moved towards the granules in a step-like manner with typically a 200-300 μm step every 20 seconds. This approach allowed the current signal to reach a pseudo-steady state at each distance travelled.

## Acknowledgements

We acknowledge the help of past group member Xue Jiang with early culture maintenance and protocol development. We acknowledge Lijang Song in the Department of Chemistry at University of Warwick for use of facilities and assistance with GC-MS analysis. We also acknowledge cooperation of Seven Trent and Draycote Water Sailing Club with sample collection. We acknowledge the help of NEOF facility members including Lucy Knowles, Rachel Tucker, Deborah Dawson and Christopher Owen, for high molecular weight DNA extractions and long-read PacBio HiFi sequencing and project design.

