## Supplementary Information for "Niche formation and metabolic interactions result in stable diversity in a spatially structured cyanobacterial community"

**SI Appendix**

**Methods and Results**

**Figures S1-S12**

**Tables S1-S6**

**References**

### Supplementary Methods and Results

#### Sample collection and culture maintenance.

Freshwater samples were collected from Draycote Water Reservoir, Warwickshire, UK on 5th October 2013. Samples were initially stored in lake water at room temperature in the laboratory (approximately 21°C) under static conditions and under diel light cycle provided by a fluorescent lamp (PowerPlant Sun Mate Grow CFL Reflector with 250w Warm Lamp).

*Irregular culture maintenance period.* Culture vessels were sealed with gas permeable film. Samples were sub-cultured in a set of media previously described for culturing of cyanobacteria: initially, samples were cultured in a minimal medium (MM) based on description in (1), consisting of a salt solution, trace metal and vitamin mixes (see Tables S4-6). Subsequently they were transferred to BG11 (2), and finally to BG11+ media (DSMZ media reference number 1593), which differs from BG11 only in vitamin B12 addition. Irregular sub-culturing was performed in liquid media or on agar plates, however, full records of culture cycles were not kept over this initial period (six years).

*Regular sub-culturing period.* Cultures were maintained in BG11+, without any carbon source addition. A vitamin mix was added to BG11+ (Table S6) and the final media used is referred to as "BG11+ vitamin mix". Cultures were grown under continuous 12h/12h light/dark cycles with white light illumination provided by a fluorescent lamp (see above). Light intensity was measured using a PAR sensor (LI-COR Quantum Sensor (LI-190R-BNC-5) and cultures were grown under 14 – 20 µmol photons $m^{-2}$ $s^{-1}$. Cultures were kept at room temperature (approximately 21°C) under static conditions and in 150 ml medical flat glass bottles. Long-term sub-culture passages have been maintained over more than two years and are ongoing. In this study, we focus on passages between 'P0' and 'P6' representing a period of one year. For each sub-culture, we performed a 1 in 200 dilution by transferring 150 µl of re-suspended filamentous culture into a final volume of 30 ml of BG11+ vitamin mix (Fig. 1A). Due to the in-homogeneity of the biofilm material, sampled biomass quantity at each transfer could not be fully standardised. Passages were performed every 34-38 days, although a time period of 187 days occurred between Passage 0 and Passage 1. Culturing conditions for the Passage 0 culture (the first sequenced passage) differed slightly from the other cultures in that the medium contained an alternative metal mixture (Table S5) to the trace metal mix of the BG11+ medium and was cultured in a 500 ml medical flat bottle with a culture volume of 100 ml. Similar filament bundles and granules were observed in this culture vessel as in later passage cultures. Regular sub-culture passages were continued beyond Passage 6 following the regime described above, for Passage 7. For Passages 8 - 10, time intervals between transfers ranged between 28 – 63 days. Light intensity and cycle were maintained but cultures were transferred to an AlgaeTron AG 230 light incubator (Photo Systems Instruments) with light supplied via cool white LEDs. From P11 to P15, dilution factor was reduced to 1:50 with a passage length of 35 days and light intensity ranged between 5 – 8 µmol photons $m^{-2}$ $s^{-1}$. Cultures were grown under fluorescent lamp illumination again from Passage 16 onwards with light intensity varying between 10 – 37 µmol photons $m^{-2}$ $s^{-1}$.

**Sub-culturing in presence and absence of vitamin mix.** A later sample (P19) of the regular passaging regime was sampled after 20 days of growth to create new cultures, using a 1:25 dilution in each of three replicate 250 ml flasks with 100 ml culture volume. A triplicate set of cultures was created for each of the BG11+ medium conditions (either with or without vitamin mix addition). Starting biomass was sampled from the parent culture by first re-suspending all biomass attached to the flask, transferring to a Falcon tube, and shaking vigorously with sterile

3 mm glass beads to disperse filamentous biofilm clumps. Cultures were incubated in the AlgaeTron AG 230 light incubator (Photo Systems Instruments) under continuous 12h/12h light/dark cycles with cool white LEDs and in-built infra-red LED lights. Light intensity ranged 10 – 14 µmol photons $m^{-2}$ $s^{-1}$. Following 35 days of culture growth, new sub-cultures were prepared by sub-culturing from each flask into a new culture flask under the same medium condition (following flask swirling and pipette mixing to disperse filaments), using a 1:50 dilution for a 30 ml culture volume in a 100 ml flask. This sub-culturing protocol was repeated three times successively, resulting in three passages from the starting cultures. After 49 days of growth per set of cultures, total biomass was harvested per flask, transferred into Falcon tubes, and freeze-dried using an Alpha 2-4 LD plus freeze-dryer (CHRIST, Germany). Three empty Falcon tubes were processed and freeze-dried alongside, to correct for weight deviation of the tubes during processing.

A linear mixed effects model was fitted to the freeze-dried biomass data using the *lme4* package (3) with R version 4.2.3 (4). Treatment (with or without vitamin mix) was used as fixed factor and Passage was used as a random factor. A likelihood ratio test was used to test significance of the fixed factor.

**Cryopreservation and revival.** We tested cryopreservation and revival of a sample taken from Passage 6 of the structured community. After 28 days of growth, filaments were re-suspended via pipette mixing and 1 ml of culture was sampled. This aliquot was preserved in 10% v/v glycerol, based on previously described protocols (5, 6). The 1 ml culture aliquot was centrifuged at 10,000 x *g* for five minutes until the culture was sufficiently pelleted, followed by removal of supernatant. 1 ml of BG11+ vitamin mix containing 10% v/v glycerol was added to the pellet and the pellet was re-suspended by pipetting. The sample was left for a 15-minute incubation period at low light intensity (below 5 µmol $m^{-2}$ $s^{-1}$). This served as equilibration to protect the cells from cryoprotectant damage. The cryotube was then stored at -80°C. After 39 days of storage at -80°C, the cryostock was revived by thawing the top of the stock so that a 300 µl aliquot could be pipetted. To wash the cells, this aliquot was added to a microcentrifuge tube then centrifuged at 6,500*g* for 5 minutes. The supernatant was discarded and fresh culture medium (BG11+ vitamin mix) without cryoprotectant was added. These centrifugations and washing steps were performed twice, and then cells were re-suspended in 300 µl of fresh BG11+ vitamin mix medium. Culture aliquot was then stored at room temperature in the dark for 24 hours. To re-grow the community culture, the 300 µl aliquot was added into 14.7 ml of BG11+ vitamin mix medium in a 50 ml Erlenmeyer flask and re-grown under the same light conditions as the original culture. A sub-culture of the revived culture was prepared after 30 days of growth, confirming longer-term culture health on re-growth.

**Samples used in taxonomy and coverage analyses.** *For short-read sequencing:* Cell pellets were collected from a set of eight samples. These eight samples consisted of *(i)* five samples taken from passages 0 and 3-6, selected for sequencing based on availability of mature granule cultures aged at 209, 254, 218, 184 and 155 days following sub-culture, respectively; *(ii)* a sample from a culture maintained in MM (see media section above), aged at 441 days since sub-culture; and *(iii)* a sample from the cryo-revived culture of passage 6 (described above) after 44 days. This last sample was compared with a similarly aged sample taken from passage 11 (collected at 49 days). An additional two samples were collected and extracted from Passages 1 and 20, at 38 and 49 days, respectively, following sub-culture. *For long-read sequencing:* Cell pellets were collected from community samples (using same methods as above) for passage 0 (aged 129 days since sub-culture) (sample 1) and passage 7 (aged 84 days since sub-culture) (sample 2), with a minimum pellet wet weight of 100 mg. For passage 0, pellets were stored at -80°C prior to DNA extraction, whereas for passage 7, pellets were snap frozen in liquid nitrogen before being stored at -80°C.

**DNA extraction and sequencing of community samples.**

Shotgun sequencing. Cell pellets were collected from samples and extracted using the Qiagen PowerSoil Pro kit (Hilden, Germany, Cat. No. 47014). Biomass (suspension or biofilm) was sampled in 1 or 1.5 ml volume for liquid culture suspension or by suction onto the end of a pipette tip for large biofilm aggregates. Tubes were centrifuged at 10,000 x *g* for five minutes in a microcentrifuge (Stuart Microfuge SCF2: Bibby Scientific, Staffordshire, UK). The liquid phase was discarded. Wet weights of the pellets were recorded and adjusted to the range of 0.04 – 0.25 grams by sampling additional culture aliquots if necessary. When pellets were not used immediately for DNA extractions, samples were stored in the -80°C freezer until extraction. For DNA extractions, the kit protocol was followed with the following modifications. Beads from each PowerBead tube provided in the kit were carefully transferred into clean microcentrifuge tubes. Cell pellets were re-suspended in 0.5 ml of sterile water then added to each PowerBead tube and centrifuged at 10,000 x *g* for five minutes. The liquid phase was removed before adding beads back into the PowerBead tubes. For the bead-beating step, a Vortex Genie-2 vortexer was used (Merck, Darmstadt, Germany, Cat No. Z258423) with a 24-tube adaptor (Qiagen, Cat No. 13000-V1-24) and all samples were vortexed for 15 minutes. For all centrifugation steps, the maximum speed of the microcentrifuge (12,300 x *g*) was set for 2 minutes. A negative control sample was included from step 1 of the protocol, by adding Solution CD1 to an empty PowerBead tube. DNA was stored in 75 µl volume of Solution C6 at -80°C before sending for sequencing. DNA concentration was quantified using a Nanodrop spectrophotometer (NanoPhotometer N60, Implen, München, Germany) and a Qubit fluorimeter (ThermoFisher Scientific, Waltham, USA, Cat No. Q33226). Total amount of raw sequence data (Gb) from Illumina shotgun NovaSeq paired-end 150 base pair read sequencing per sample was as follows: P0 = 10.8, P3 = 15.3, P4 = 14.4, P5 = 14.7, P6 = 15.2.

PacBio HiFi sequencing. DNA extractions were performed by the Natural Environment Research Council (NERC) Environmental Omics Facility (NEOF). In brief, high molecular weight (HMW) DNA was extracted using the Macherey-Nagel NuceloBond HMW DNA kit with liquid nitrogen grinding using mortar and pestle for more than 15 minutes. The frozen culture sample was added directly to the kit buffer in a 50 ml tube to reduce any HMW DNA degradation. Proteinase K volume was doubled, compared to kit instructions. The samples were then cleaned using AMpure PB beads with four repeated cycles. DNA was extracted in small volumes to reduce contaminants and DNA degradation. DNA quality scores were 1.2 – 1.5, measured by absorbance at 260/230nm and 1.75 – 2.0 measured by absorbance at 260/280nm. PacBio DNA libraries and sequencing were completed at the Centre for Genomics Research (CGR), from extracted genomic DNA samples. Following sample QC, low input library preparation was used for sample 1 (sample P0) and the ultra-low input protocol was used for sample 2 (sample P7). Sequencing was performed on the Sequel II SMRT Cell in CCS run mode and raw sequence data was delivered for downstream bioinformatics analyses.

**Long-read sequence assembly and binning.** The two PacBio HiFi samples (described above) were assembled using hifiasm-meta (7) and the resulting unitig assembly graphs were used in the downstream analyses. ORFs were called on unitigs with Prodigal (V2.6.3) (8) with option meta, and single-copy core genes (SCGs) were annotated through RPS-BLAST (v2.9.0) (9) using the pssm formatted COG database (10), which is made available by the CDD (11) as in the STRONG pipeline. However, to take into account strain diversity and non bluntification of the assembly graph, SCGs were clustered with MMseqs2 (12) (v13.45111) with options --minseq-id 0.99 -c 0.80 --cov-mode 2 --max-seqs 10000. Reads were mapped to the assembly with minimap2 (2.17-r974-dirty) (13) using preset -ax asm10. Coverage was obtained from alignment using samtools (1.17) (14) and bedtools (v2.25.0) (15). The unitigs were then binned from their coverage and composition, using both binning software CONCOCT (16) and metabat2 (17), respectively allowing unitigs as small as 1000bp and 1500bp. The two resulting sets of MAGs were combined in a unique set using custom scripts resulting in 18 high quality bins (greater than 75% completeness of SCGs in single-copy). For two high abundance species, *F. draycotensis* and the species from the *Chryseoglobus* genus- for which the genome was split over three bins- two separate complex (high variability) circular components were observed in the assembly graph. In these cases, we found circular consensus paths using maximum aggregate coverage depths. Of the resulting 16 long-read genomes (MAGs), 5 were contained in a single, circular contig indicating high quality assemblies, whereas 11 were split over multiple contigs (see Table S3). We taxonomically classified MAGs with GTDB-Tk v2.1.0 (18) and data version r207, using standard settings on GTDB-Tk, revealing that two of these were strains of *Allorhizobium rhizophilum*. For the MAG identified as deriving from the *Phreatobacter* genus, genome size was small and completeness based on Check M using the DFAST annotation platform (19) (see below) was only 48.36 %. We improved on this by using the connected graph component for this MAG instead (Comp 243), which had higher completeness (see Table S3).

To assess the fraction of the total diversity represented by this collection of MAGs, 16S genes were annotated in the assembly using Barrnap (20) and clustered into OTUs using VSEARCH (21) with 97% identity. Each of the resulting 14 distinct OTUs (species and strains of the *Allorhizobium* genus were clustered together) could be mapped to one or more MAGs, which allows to ascertain that the full genomic diversity in the dataset was converted into MAGs.

**Genome annotations for metabolic pathways**. Genome annotations were performed using the DFAST annotation platform (releases 1.2.15/1.2.18; (19)). DFAST was run per genome with settings additional to the defaults as follows: perform taxonomy and completeness checks, use Prodigal for annotation of the coding sequence, set E-value to 1e-10 and enable both HMM scan and RPSBLAST. For the cyanobacterial genome, the Cyanobase organism-specific database was selected. Resulting protein sequences from genome annotation were further annotated for KEGG orthologs (KOs) using KofamKOALA (release 102.0/103.0; (22)). Resulting KO lists from short and long-read data were concatenated to create a unique KO list (where possible) for each of the 17 species in the final set. KO lists were combined for the two bins predicted to represent two different strains of *Allorhizobium rhizophilum* in the long-read data. Metabolic module completeness based on KOs was completed using the MetQy tool (23). For module sets associated with carbon, amino acid, cofactor and vitamin metabolism on the KEGG database (24), a threshold for completeness at 0.25 across genomes for a given module was applied (Fig. S4, 7-8). Individual KOs were manually searched for the vitamin B7, B12 and sulfur pathways analyses (*Supplementary files* 3 - 5). Data was plotted using R version 4.1.2 (25) and package MetQy (version 1.1.0) (23).

**Analysis of vitamin biosynthesis pathways**. Vitamin B7 (biotin) biosynthesis consists of a two-part pathway (illustrated in Fig. 4A) (26) and the genes *bio ABDF* of the lower pathway were mostly absent across all species apart from *P. composti* (Fig. 4A and *Supplementary file* 3). The vitamin B12 (cobalamin) biosynthesis pathway consists of more than 30 genes, involving corrin ring synthesis and the nucleotide loop assembly (27, 28) (Fig. S9B and D and *Supplementary file* 4). Between these two pathways, several genes are overlapping (Fig S9A). Additionally, synthesis of the ligand DMB (5,6-dimethylbenzimidazole) from riboflavin (29,

30) (Fig. S9D) is required, defined on the KEGG database (24) and in (29). Capability for riboflavin biosynthesis is present across several genomes (Fig. S9). Cells can alternatively transport corrinoid compounds (such as the intermediate cobinamide) into the cell and convert the intermediate cobinamide into cobalamin via a scavenging pathway (28) (Fig S9D). This pathway shares overlapping genes with those of the nucleotide loop assembly. Some transporters for cobalamin import have been differentially characterised for gram-negative and positive bacteria (28, 31) (Fig S9C) however the presence of these was mainly incomplete across our species. In our system, we found that *P. composti* and *A. rhizophilum* have complete or near-complete modules for all sections of the vitamin B12 pathway (Fig. S9A). All other species have partially complete B12 pathways, some also missing DMB synthesis but with cobinamide scavenging complete, including *F. draycotensis* (Fig. S9A). Vitamin B12 biosynthetic genes are almost completely absent in the species from the *Microbacteriaceae* family and in the three species from the *Acetobacteraceae* family. Vitamin B12 biosynthesis is so far known to be restricted to certain bacterial species (32, 33) however species variation in genes encoding biosynthesis and transport pathways is under-characterised. The alternative Gram-negative transporter BtuM that can substitute for BtuCDF (28) does not have an assigned KO in the KEGG database, so its presence was not analysed.

**Phylogenetic analyses.** Taxonomic assignment of metagenomes and circularised genomes was performed using GTDB-Tk v2.1.1 (18) and data version r207, using standard settings on GTDB-Tk. This approach involves placing user MAGs on a pre-compiled phylogenetic tree of existing species and MAGs found on GTDB (34). The resulting tree contains over 3600 tips, which represent the GTDB database species (or MAGs) and the MAGs from our sample. To facilitate its visualisation, in Fig. 1C, we have randomly sampled 5 percent or 1 representative (whichever was larger) of GTDB database tips at the order-level, except for orders that had 10 or fewer representatives, which were included with all their representatives. This process resulted in 2590 species (or MAGs), including the 16 MAGs discovered in this study. Out of these 16 MAGs, 12 show high similarity to cultured or sequenced genomes and were assigned by GTDB-Tk at genus level, one was assigned at family level whilst the other three were assigned at species level (see Table 1).

In the case of the cyanobacterium found in the presented community, there was high sequence similarity to only one other uncultured metagenome in the databases (GTDB id; JAAUUE01 sp012031635). This prompted us to further explore the taxonomic placement and run an additional phylogenetic analysis. To do so, we used the multiple sequence alignment of the single copy core genes (SCGs) – as created by the GTDB-Tk platform – to create an alignment of the cyanobacterium MAG identified in this study, all the GTDB species/MAGs from the Cyanobacteriales order, and the *Pseudomonas_E composti* MAG identified in this study (as an outgroup). We then used this alignment to build a maximum likelihood tree with FastTree (35) using the default options. The resulting tree, re-rooted at the *P. composti* MAG, is shown in Fig. S2.

**Species isolation.** Isolation of species from the community was achieved using three types of agar with carbon source supplementation: BG11+ vitamin mix with 0.1% w/v glucose, yeast mannitol and BG11+ vitamin mix with 0.05% w/v riboflavin. Yeast mannitol was selected to enrich for *A. rhizophilum* and was prepared following the methods of (36). Riboflavin was chosen to select for *M. maritypicum* based on previous description of riboflavin degradation by this species (37). Addition of glucose was expected to generally enrich for heterotrophs.

Later passage cultures were selected to attempt isolation, using pipette mixing to resuspend filaments. For BG11+ vitamin mix with 0.1% w/v glucose, a 10 µl aliquot from passage 8, aged 42 days old, was sampled and successively streaked across an agar plate.

Additionally, a later passage culture was diluted into BG11+ vitamin mix with 0.1% w/v glucose medium and subsequently aliquoted and streaked onto agar plates of the same medium and LB plates. For yeast mannitol, a dilution of 1000-fold was first prepared from passage 7, aged 60 days old. A spread plate was prepared with a 50 µl aliquot from this dilution. For BG11+ vitamin mix with 0.05% w/v riboflavin, a 100 µl aliquot was sampled from passage 12 at 57 days old and a spread plate was prepared. Agar plates were either incubated under the same light conditions as the community culture (see above) for BG11+ vitamin mix + 0.1% glucose, or in the dark at 30°C for 2 – 3 days, or up to 9 days for BG11+ vitamin mix with riboflavin due to slower colony growth. Single colonies observed on each agar type were re-streaked onto new agar plates of the same type either once more (riboflavin agar), three times (glucose agar) or four times (yeast mannitol agar) until maintenance of the colony morphotype was clear. Plates were incubated at 30°C in the dark. On the glucose plate, a cream-coloured smooth margin colony type was observed. On the yeast mannitol plate, round cream colonies were observed with a gelatinous texture. On the riboflavin plate, small, cream, round colonies were visible and bleaching of the plate occurred due to riboflavin breakdown (38).

A liquid culture of each colony type was prepared to harvest material for DNA sequencing and for cryopreservation. A single colony was inoculated into 30 ml of each respective medium and grown in a 100 ml Erlenmeyer flask, incubated at 30°C and 180 rpm for 48 hours. For the species isolated on BG11+ vitamin mix with 0.05% w/v riboflavin, the liquid culture was prepared in LB medium as growth was unsuccessful in liquid riboflavin medium. A liquid LB culture was also prepared for the species isolated on BG11+ vitamin mix + 0.1% glucose and LB agar plates. Cryostocks were prepared with a final concentration of 15% v/v glycerol for the cultures grown in BG11+ media and with 30% v/v glycerol (as of (39)) for the culture grown in yeast mannitol medium. Cryostocks were stored at -80°C. The remaining cultures were pelleted at 4000 rpm for 10 minutes. DNA was extracted from the cell pellet using the Qiagen PowerBiofilm kit (Cat. No. 24000-50) and stored at -80°C. DNA concentration was quantified using the Nanodrop spectrophotometer.

**Species characterisation via 16S rRNA gene region amplification.** Isolates were characterised via Sanger sequencing of the bacterial 16S rRNA V3-V4 gene region using primers 341F and 806R as described in (40). To distinguish between the two species of the *Allorhizobium* genus, a longer region was amplified using primer 341F with 1391R (41) as sequence differences occurred between the two species in this region. DNA was diluted to a concentration of approximately 10 ng/µl and added in 5 µl volume to the PCR reaction mix of 25 µl final volume. The reaction mix consisted of 12.5 µl of GoTaq G2 Green Master Mix (2X) (Promega Product Code: M7822), 1 µl of each of the forward and reverse primers (10 µM concentration) and 20 µl of sterile MilliQ water. For some PCR runs, a total volume of 50 µl was used and in these cases, the reagent volumes were doubled.

The PCR reaction was run using an Applied Biosystems Veriti Thermal Cycler (California, USA) with the following cycling conditions. Firstly, an initial denaturation of three minutes at 95°C was run. This was followed by 30 cycles consisting of denaturation for thirty seconds at 95°C, annealing at 49°C (50°C for 341F/1391R) for thirty seconds and elongation for ninety seconds (sixty seconds for 341F/1391R) at 72°C. A final elongation step of ten minutes (seven minutes for 341F/1391R) at 72°C was run followed by infinite hold at 4°C.

Products were run on a 1% w/v agarose gel stained with GelRed dye (Biotium, product code: 41003) using 1 x TAE buffer, including a 100 bp ladder (NEB, product code: N3231S) mixed with purple loading dye. The gel was run at 100 V for at least 45 minutes. Bands were visualised with the gel imaging system U:Genius3 (Syngene, Cambridge, UK) via a blue LED transilluminator. A band of the correct size could be visualised between 400 and 500 base pairs for primer pair 341F/806R and around 1 kilo base pair for primer pair 341F/1391R. PCR products were purified using the GeneJet gel extraction kit (Thermo Scientific, K0691) with the following modifications. Centrifugations were performed at 12, 300 x *g* for 60 seconds. Sodium acetate was not added. Purified products were eluted in 20 – 50 µl of elution buffer then DNA concentration was quantified on the Nanodrop spectrophotometer. DNA was Sanger-sequenced (GATC Sequencing Service) in the forward direction using the same forward primer as in the PCR reaction. Taxonomic identity was characterised using sequence alignment to the annotated 16S rRNA gene sequences from the long-read shotgun metagenomics community data.

Characterisation of the 16S rRNA gene regions distinguished the following species: *P. composti*, *A. rhizophilum*, *A. sp900156055* and *M. maritypicum*. These species were grown successfully on BG11+ vitamin mix with 0.1% w/v glucose, yeast mannitol and BG11+ vitamin mix with 0.05% w/v riboflavin, respectively.

**Isolated species physiological assays: Biolog carbon sources.** To assess growth of the isolated species on different carbon sources, the Biolog phenotypic microassay 'PM1 96 Carbon Utilisation Assay' (Biolog, Hayward, CA, USA) was used. This assay includes a range of possible carbon sources, including a set of amino acids, nucleotides, organic acids, polymers, sugars, sugar alcohols and sugar phosphates (42). Each species was revived from its cryostock on LB agar. Plates were incubated at 30°C for 24 hours (for *P. composti*) and for 48 hours for the other species (until sufficient colonies had grown). Each species was re-streaked onto a new LB agar plate before use in the Biolog assay. The inoculating fluid 1.0 x IF-0a to be used to suspend colonies swabbed from the agar plate, was prepared from 1.2 x IF-0a (Technopath, UK, product code: 72268) by dilution. For the Gram-positive protocol, an additive solution was added to the inoculating fluid consisting of the following components in their final concentrations: 2mM magnesium chloride hexahydrate, 1mM calcium chloride dihydrate, 25µM L-arginine HCl, 50µM L-glutamate Na, 12.5µM L-cystine pH8.5, 25µM 5'-UMP 2Na, 0.005% yeast extract, 0.005% tween-85.

Colonies were removed from the LB agar plate using a sterile swab and added to IF-0a to form a lightly turbid suspension. The transmittance (%T) at 600 nm wavelength (turbidity) was checked using a benchtop spectrophotometer (Spectronic 200E, Thermo Scientific), and culture density was adjusted until %T was approximately 42% or 81% for the gram-negative (43) and gram-positive (44) species respectively. For gram-negative species, cell suspension was then diluted in 1.0 x IF-0 to the starting inoculum of 85% T (+/- 3%), equivalent to 0.07 OD (43). For gram-positive species, cell suspension was further diluted in inoculating fluid and additive solution to the starting inoculum of 98% T (+/- 2%), equivalent to 0.007 OD. Plate PM1 was inoculated with 100 µl per well of cell suspension, with a separate plate per species.

Growth was measured via absorbance (optical density at 600 nm wavelength) with continuous incubation in a plate reader (CLARIOstar, BMG LABTECH GmbH, Ortenberg, Germany) at 30°C for 48 hours. Measurements were taken every fifteen minutes with double orbital shaking at 200 rpm for five minutes before each reading. Endpoint readings (48 hours) were used for data analysis, blank-corrected by the first or second optical density reading per well. Data was presented using R version 4.1.2 (25).

**Isolated species physiological assays: growth in the presence or absence of vitamin mix.** To assess growth in BG11+ medium with vitamin mix removal, we tested growth of isolated species *P. composti* and *A. rhizophilum.* Firstly, BG11+ medium was prepared as a base stock and supplemented with 0.8% w/v glucose to enable growth, then split into two media: one with vitamin mix added and one with water added as replacement. An aliquot of cryostock of each species was revived by streaking to single colonies on LB agar and incubating at 30°C for up to 48 hours. An overnight culture per species was prepared by picking an individual colony and inoculating into liquid LB medium for up to 24 hours at 30°C and 169 rpm. Cultures were then diluted to $OD_{600nm}$ = 1.0 on a benchtop spectrophotometer, before washing twice and re-suspending in 0.9% w/v saline at equal volume. Each culture was split and re-suspended in each of the medium types. Culture was added in three (*P. composti*) or four (*A. rhizophilum*) replicate wells in 200 µl final volumes in a Greiner flat-bottomed non-treated 96-well plate per species (Catalog no: 655161), performing a 1:100 dilution into the respective medium for a starting density of approximately $OD_{600nm}$ = 0.01. Growth was measured in the plate reader (CLARIOstar, BMG LABTECH GmbH, Ortenberg, Germany) following the settings described in the section above, for 48 hours. A path length correction of 1 cm was applied for 200 µl well volumes. Growth assays were performed on separate days for the two species.

For *A. rhizophilum*, growth was performed in BG11+ medium supplemented with 0.8% w/v glucose, with two additional medium conditions. For one condition, BG11+ medium was supplemented only with biotin at the same concentration as in the vitamin mix. For the other medium condition, supernatant from a grown culture of *P. composti* was mixed with BG11+ medium without vitamin supplementation (1.25 x reagents), in a 20:80 ratio. Supernatant was prepared as follows. A large culture of *P. composti* was grown from inoculum prepared for the 96-well plate assay described above. 200 µl of cell suspension adjusted to $OD_{600nm}$ = 1.0 was inoculated in a final volume of 20 ml of BG11+ medium without vitamin mix for a 1:100 dilution and grown in a 50 ml falcon tube for 48 hours at 30°C and 169 rpm. Following growth, an $OD_{600nm}$ of 0.819 was measured in a benchtop spectrophotometer. Culture supernatant was filter-sterilised through a 0.22 µm syringe filter unit and stored in the fridge until use.

**Bacterial imaging of community sample**

A 1 mL sample of culture (32 days after 1:50 subculture) was centrifuged at 4000 rpm for 5 minutes to pellet cells, which were then resuspended in PBS with 2% DMSO solution, containing a final concentration of 10 µM Thioflavin T (ThT). The sample was allowed to incubate for 20 minutes at room temperature. For post-stain washing, the sample was centrifuged at 4000 rpm for 5 minutes, and the pellet resuspended in fresh BG11+ vitamin mix medium in the same volume. 30 µl of stained sample was added to a BG11+ vitamin mix agar pad, allowed 1-2 minutes to settle and then inverted into a cover-glass bottom dish (WillCo Wells, USA, HBST-5040). The sample was incubated for 30 minutes on the bench, before observing on an Olympus IX83 inverted fluorescence microscope with 10x objective (Olympus UPLFLN10X2PH), excited with the CoolLED pE-300 light system. ThT was observed through a ECFP cube (U-F9001), whilst the fluorescence from the cyanobacterial photopigment was observed through a HcRed cube (U-F41043). Note that ThT is known to enter bacterial cells depending on their membrane potential, and indicate their active, live state (45). Fig S5 is representative of replicate samples on agar pads, showing a greater concentration of bacteria within the slime tracks than evenly distributed across the agar pad.

**Slime extraction for GC-MS characterisation**

Slime extraction was performed adapting the methods of (46) and (47). The final protocol is available via protocols.io (48). In brief, the pellet underwent hydrolysis with sulfuric acid, and then trimethylsilylation in preparation for the GC-MS. GC-MS was performed on an Agilent 7890GC, coupled with a 5977B MSD detector. An Agilent HP-5MS with 5% Phenyl Methyl Silox column was used (30 m × 250 µm × 0.25 µm). The initial column temperature was set to 150ºC, held for 2 minutes, and increased at a rate of 8ºC/min to 250ºC, then maintained for 17.5 minutes. Helium was used as a carrier gas (1.2 mL/min). The front inlet temperature was 275°C, the transfer line temperature was 280°C, the MS source temperature was 230°C, and the MS quad temperature was 150°C. An injection volume of 1 µL was used, with an injection dispense speed of 6000 μL/min. The total run time was 32 minutes. Electron ionisation was used, with a MS scan range 50-750 m/z.

As part of the GC-MS work on monosaccharide composition of the slime/EPS material, the mass spectra can be examined to confirm a spectral match between the standards and the slime. Fig. S6 shows different mass spectra collected at different time points of the GC spectra. Many of the monosaccharides have the same molecular mass and very similar structure, which when fractionated produce identical fragments. This is seen in the similarities between the mass spectra for the glucose and galactose standards (Fig. S6A and C, respectively). These fragment patterns match well to those seen in the slime, observed at the peaks which best match the retention times. Whilst the limited specificity of the MS data cannot absolutely confirm the identity of the monosaccharides, the similar fragment patterns combined with the matching peak retention time give confidence in identifying galactose and glucose in the slime.

**Micron-scale probe measurement of oxygen and hydrogen sulfide gradients**

Oxygen was measured using a Unisense OX-NP-710704 probe, whilst hydrogen sulfide was measured using a Unisense H2S-500-305717 probe. Probes were calibrated on the day of measurement, using kits provided by Unisense. The calibration kit for oxygen measurement was a Unisense zero-oxygen calibration kit (Product number: Calkit-O2) and for hydrogen sulfide measurement the kit used was a Unisense H2S/Sulf Sensor calibration kit (Product: Calkit-H2S). Granules were sampled across a set of different cyanobacterial cultures of different passage numbers and ages (Fig. 3D and S12). Data is only shown for oxygen measurements, as the hydrogen sulfide profiles were not consistent across granules. For Fig. 3D, a culture equivalent to Passage 23 was sampled. Cultures sampled for Figure S12 are indicated in the legend.

### Supplementary Figures

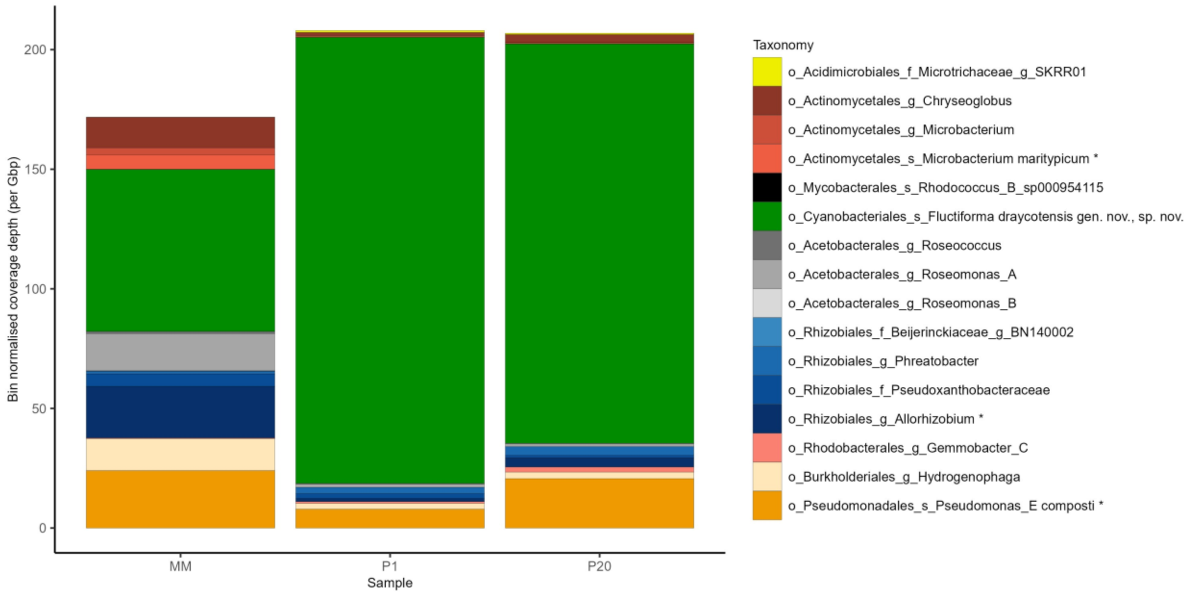


**Figure S1. Coverage of taxonomic bins in additional community samples.** Samples shown are as follows: MM = early sample grown in minimal medium (MM) from ‘irregular passaging' period aged 441 days; P1 = Passage 1 sample aged 38 days; P20 = Passage 20 sample aged 49 days. Samples P1 and P20 were not co-assembled with the other passage samples and MM, and instead analysed separately, mapping reads onto the MAGs identified from the other samples. Community composition, i.e. species coverage, colour-grouped at the taxonomic order level (listed in Table 1 and shown on the legend) for 16 species characterised via short-read sequencing. Colour shades indicate MAGs within orders and the highest taxonomic resolution is described in the legend on the figure. 'o' = order; 'f' = family; 'g' = genus; 's' = species. Asterisks indicate species that were isolated from later passage community cultures. See *Methods* for sample details. The y-axis shows bin normalised coverage depth per Gbp of sequencing.

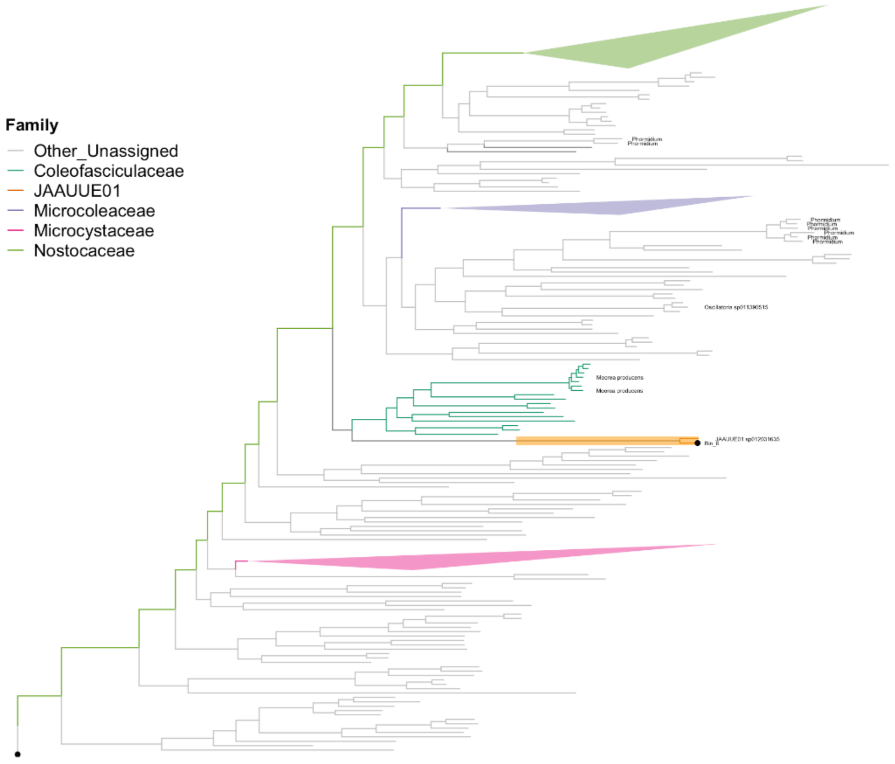


**Figure S2. Maximum likelihood phylogenetic tree for the cyanobacterial species identified in this study.** See *Methods* for tree creation. Species included are known species of the Cyanobacteriales order obtained from GTDB, the single new cyanobacterial species identified in this study (labelled as Bin 6 on the tree), and the *P. composti* bin from this study. Both of these bins are indicated with a black dot on the tree. The tree is re-rooted at the *P. composti* bin, which was included as an outgroup. The branch length to this bin was divided by 50 to better visualise the tree. Key families of the Cyanobacteriales order are highlighted as indicated in the legend, with the *Nostocaceae* (183 species), *Microcoleaceae* (45 species), and *Microcystaceae* (41 species) family clades collapsed to better visualise the tree. Notice that the commonly studied filamentous cyanobacterial genera *Nostoc* and *Trichodesmium* fall under the *Nostocaceae* and *Microcoleaceae* families respectively. Notice that several species named under the *Oscillatoria* and *Phormidium* genera, which have also been studied in the past for their motility, do not form a monophyletic group. These species, as well as several species of the genus *Moorea*, studied for their natural product biosynthesis (49), are highlighted on the tree. The JAAUUE01 family is automatically generated by the GTDB database and contains only two species, the one discovered and cultured in this study, and another uncultured MAG (JAAUUE01 sp012031635) (50).

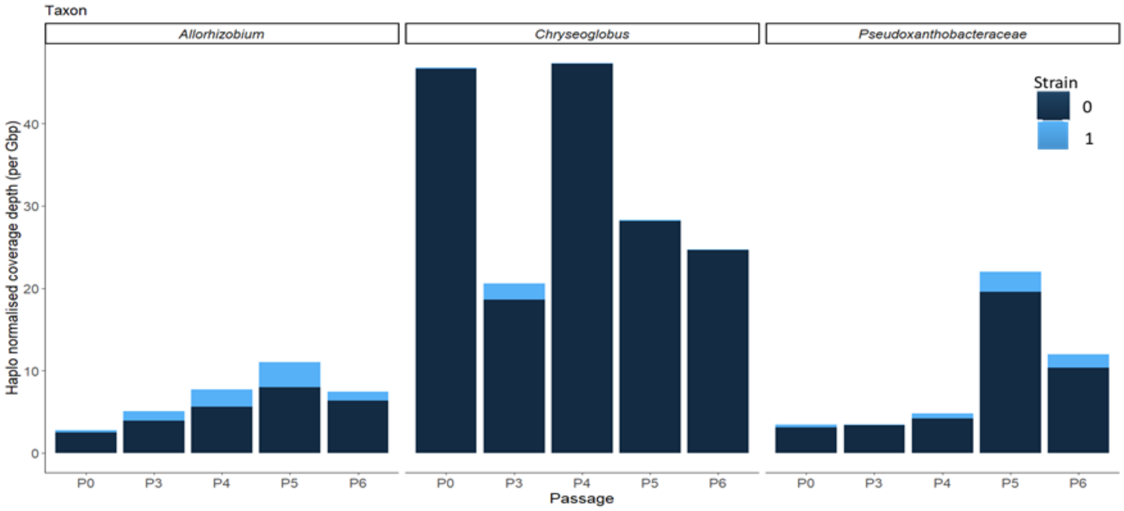


**Figure S3. Within-bin haplotype coverage over passaged cultures.** Coverage was calculated as an intensity value with the Bayes path algorithm, calculated as k-mer coverage across single copy genes (SCGs) divided by read length within the STRONG pipeline (see *Methods*). The intensity proportion assigned to each haplotype per bin was multiplied by the normalised coverage depth per Gbp for the bin for each passage, to calculate haplotype normalised coverage depth. Two strains were detected in each of three bins (species) indicated in the header of each sub-plot (see Table 1), over the five presented passages.

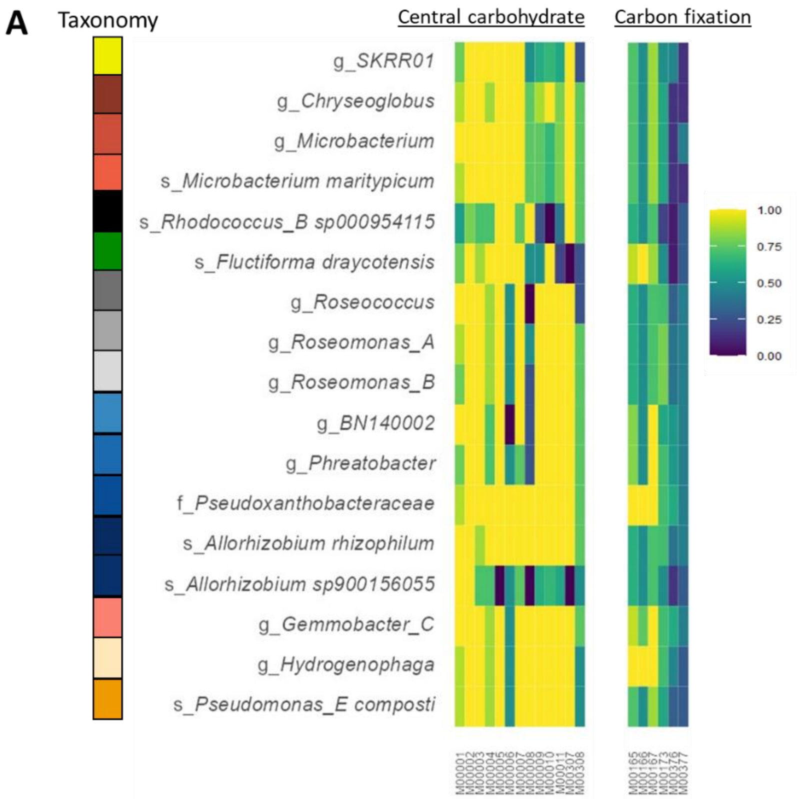


**B**

| Central carbohydrate metabolism | |
|---|---|
| M00001 | Glycolysis: glucose =>pyruvate |
| M00002 | Glycolysis_core |
| M00003 | Gluconeogenesis |
| M00004 | Pentose Phosphate Pathway |
| M00005 | PRPP biosynthesis, ribose 5P => PRPP |
| M00006 | PP_ox, glucose 6P => ribulose 5P |
| M00007 | PP_nonOx, fructose 6P => ribose 5P |
| M00008 | ED, glucose-6P => glyceraldehyde-3P + pyruvate |
| M00009 | TCA |
| M00010 | TCA_1, oxaloacetate => 2-oxoglutarate |
| M00011 | TCA_2, 2-oxoglutarate => oxaloacetate |
| M00307 | Pyruvate oxidation, pyruvate => acetyl-CoA |
| M00308 | ED_1, gluconate => glycerate-3P |

| Carbon fixation | |
|---|---|
| M00165 | Reductive PP (Calvin cycle) |
| M00166 | Reductive PP_1, ribulose-5P => glyceraldehyde-3P |
| M00167 | Reductive PP_2, glyceraldehyde-3P => ribulose-5P |
| M00173 | Reductive citrate cycle (A-B) |
| M00376 | 3-Hydroxypropionate bi-cycle |
| M00377 | Reductive acetylCoA (WL) |

**Figure S4. Completeness of KEGG metabolic modules associated with carbohydrate metabolism. (A)** Selected modules for central carbohydrate metabolism and carbon fixation as defined on the KEGG database (24) and analysed using the MetQy tool (23) (see *Methods*). Carbon fixation modules were selected that apply to bacterial fixation pathways. Only modules that were present to a level of completeness above 0.25 in at least one MAG are shown, combining annotation from short-read and long-read genomes. Bins are colour-categorised by order (as in Fig. 2) and named by the highest taxonomic resolution resolved per bin (family, genus or species). Full taxonomy is given in Table 1. **(B)** Module definitions from KEGG database (24). Abbreviations are as follows: PRPP = Phosphoribosylpyrophosphate, ED = Entner-Doudoroff pathway, A-B = Arnon-Buchanan cycle, WL = Wood-Ljungdahl pathway. Three alternative carbon fixation pathways were analysed: reductive citrate cycle (A-B): M00173, reductive acetylCoA (WL): M00377 and 3-hydroxypropionate bi-cycle: M00376.

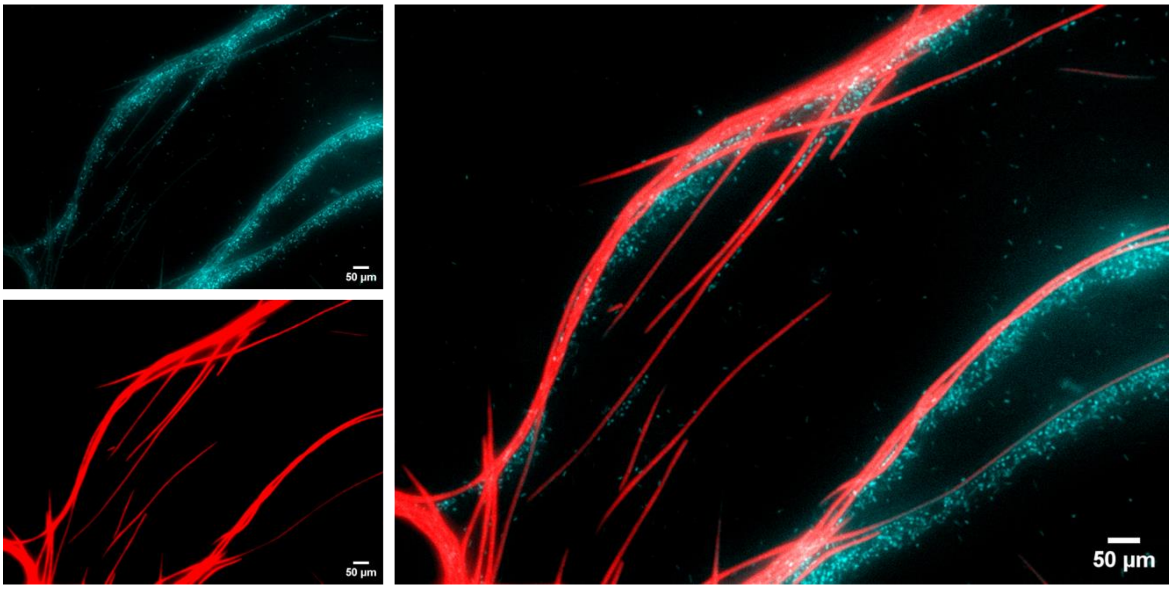


**Figure S5. Community sample stained with 10 µM Thioflavin T (ThT).** Left panels show fluorescence across different filters, with the ThT fluorescence measured through a ECFP (U-F9001) filter cube (top, false coloured in cyan), and the cyanobacterial photopigment fluorescence recorded through a HcRed (U-F41043) filter cube (bottom, false coloured in red). The right panel shows the combined image. Cells were placed on a BG11+ vitamin mix medium agar pad, which was then flipped onto a cover slide (see SI methods). Note that ThT is known to be taken up by bacteria and indicates their membrane potential and active live state (45).

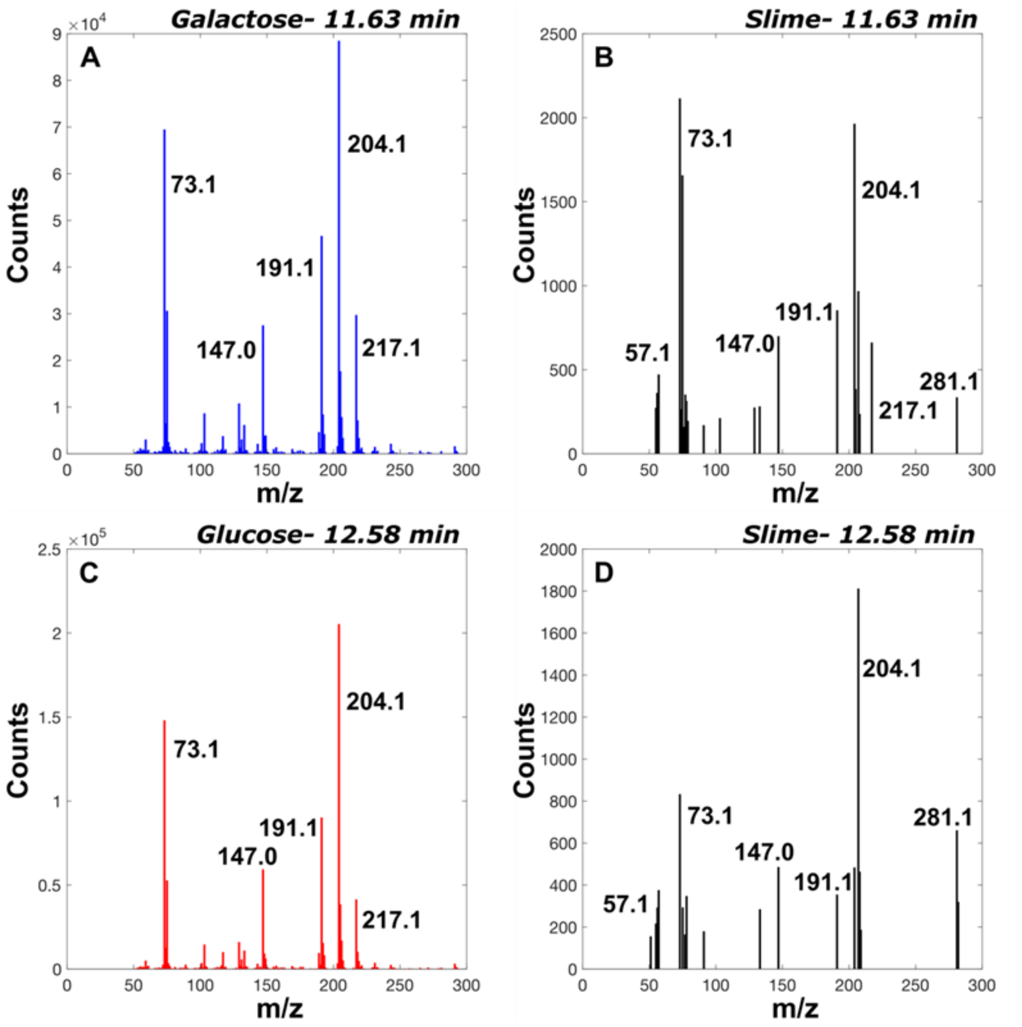


**Figure S6. Accompanying mass spectra for the GC analysis shown in Figure 3C.** Mass spectra at the 11.63 min peak are shown for the galactose standard (A) and for the corresponding peak in the slime GC (B). Mass spectra at the 12.58 min peak are shown for the glucose standard (C) and for the corresponding peak in the slime GC (D).

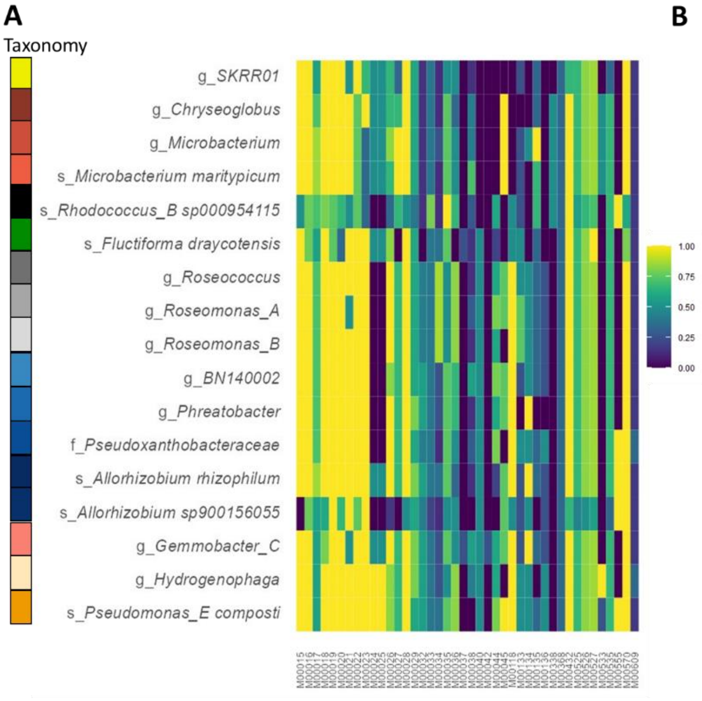


B

| Amino acid metabolism | |
|---|---|
| M00015 | Proline |
| M00016 | Lysine |
| M00017 | Methionine |
| M00018 | Threonine |
| M00019 | Valine |
| M00020 | Serine |
| M00021 | Cystine |
| M00022 | Shikimate |
| M00023 | Tryptophan |
| M00024 | Phenylalanine |
| M00025 | Tyrosine 1 |
| M00026 | Histidine |
| M00027 | GABA 3 |
| M00028 | Ornithine |
| M00029 | Urea_cycle |
| M00032 | Lysine deg |
| M00033 | Ectoine |
| M00034 | Methionine salvage |
| M00035 | Methionine deg |
| M00036 | Leucine deg |
| M00037 | Melatonin |
| M00038 | Tryptophan metab |
| M00040 | Tyrosine 2 |
| M00042 | Catecholamine |
| M00044 | Tyrosine deg |
| M00045 | Histidine deg |
| M00118 | Glutathione |
| M00133 | Polyamine |
| M00134 | Polyamine 2 |
| M00135 | GABA |
| M00136 | GABA 2 |
| M00338 | Cystine 1 |
| M00368 | Ethylene |
| M00432 | Leucine |
| M00525 | Lysine 2 |
| M00526 | Lysine 3 |
| M00527 | Lysine 4 |
| M00533 | Homoprotocatechuate |
| M00535 | Isoleucine |
| M00555 | Betaine |
| M00570 | Isoleucine 2 |
| M00609 | Cystine 2 |

**Figure S7. Completeness of KEGG metabolic modules associated with amino acid metabolism.** (A) Select modules as defined on the KEGG database (24) and analysed using the MetQy tool (23) (see *Methods*). Only modules that were present to a level of completeness above 0.25 in at least one MAG are shown, combining annotation from short-read and long-read genomes. Bins are colour-categorised by order (as in Fig. 2) and named by the highest taxonomic resolution resolved per bin (family, genus or species). Full taxonomy is given in Table 1. (B) Module definitions from KEGG database (24).

**A**

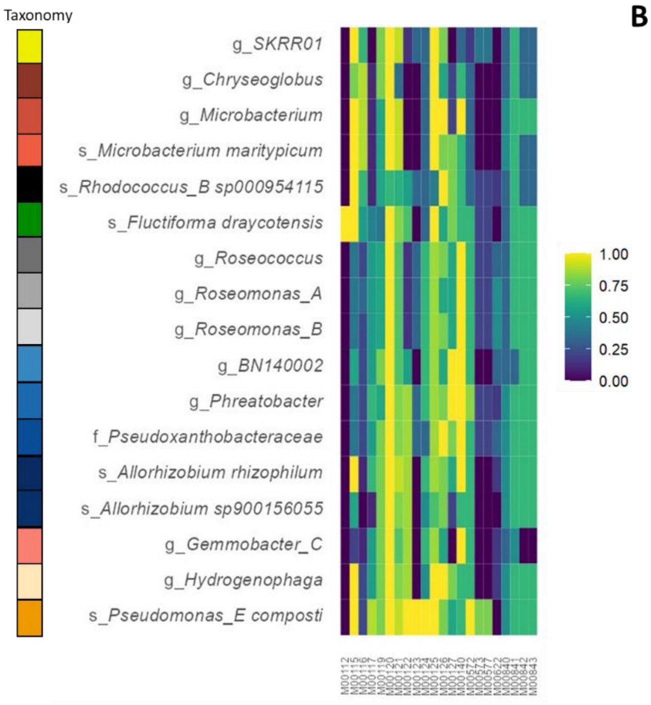


**B**

| Vitamin and cofactor biosynthesis | |
|---|---|
| **M00112** | **Tocopherol/tocotorienol** |
| M00115 | NAD |
| M00116 | Menaquinone |
| M00117 | Ubiquinone |
| M00119 | Pantothenate (vitamin B5) |
| M00120 | Coenzyme A |
| M00121 | Heme |
| **M00122** | **Cobalamin, cobyrinate a,c-diamide => cobalamin (vitamin B12)** |
| **M00123** | **Biotin, pimeloyl-ACP/CoA => biotin (vitamin B7)** |
| **M00124** | **Pyridoxal-P (vitamin B6)** |
| M00125 | Riboflavin (vitamin B2) |
| M00126 | Tetrahydrofolate (vitamin B9) |
| M00127 | Thiamine (vitamin B1) |
| M00140 | C1-unit interconversion |
| **M00572** | **Pimeloyl-ACP, BioC-BioH pathway (vitamin B7)** |
| M00573 | Biotin, BioI pathway, long-chain-acyl-ACP => pimeloyl-ACP => biotin |
| M00577 | Biotin, BioW pathway, pimelate => pimeloyl-CoA => biotin |
| M00622 | Nicotinate degradation (vitamin B3) |
| M00840 | Tetrahydrofolate, mediated by ribA and trpF (vitamin B9) |
| M00841 | Tetrahydrofolate, mediated by PTPS (vitamin B9) |
| M00842 | Tetrahydrobiopterin |
| M00843 | L-threo-Tetrahydrobiopterin |

**Figure S8. Completeness of KEGG metabolic modules associated with metabolism of cofactors and vitamins. (A)** Select modules as defined on the KEGG database (24) and analysed using the MetQy tool (23) (see *Methods*). Only modules that were present to a level of completeness above 0.25 in at least one MAG are shown, combining annotation from short-read and long-read genomes. Bins are colour-categorised by order (as in Fig. 2) and named by the highest taxonomic resolution resolved per bin (family, genus or species). Full taxonomy is given in Table 1. **(B)** Module definitions from KEGG database (24). Those highlighted in bold indicate full module completeness in just a single species of the community.

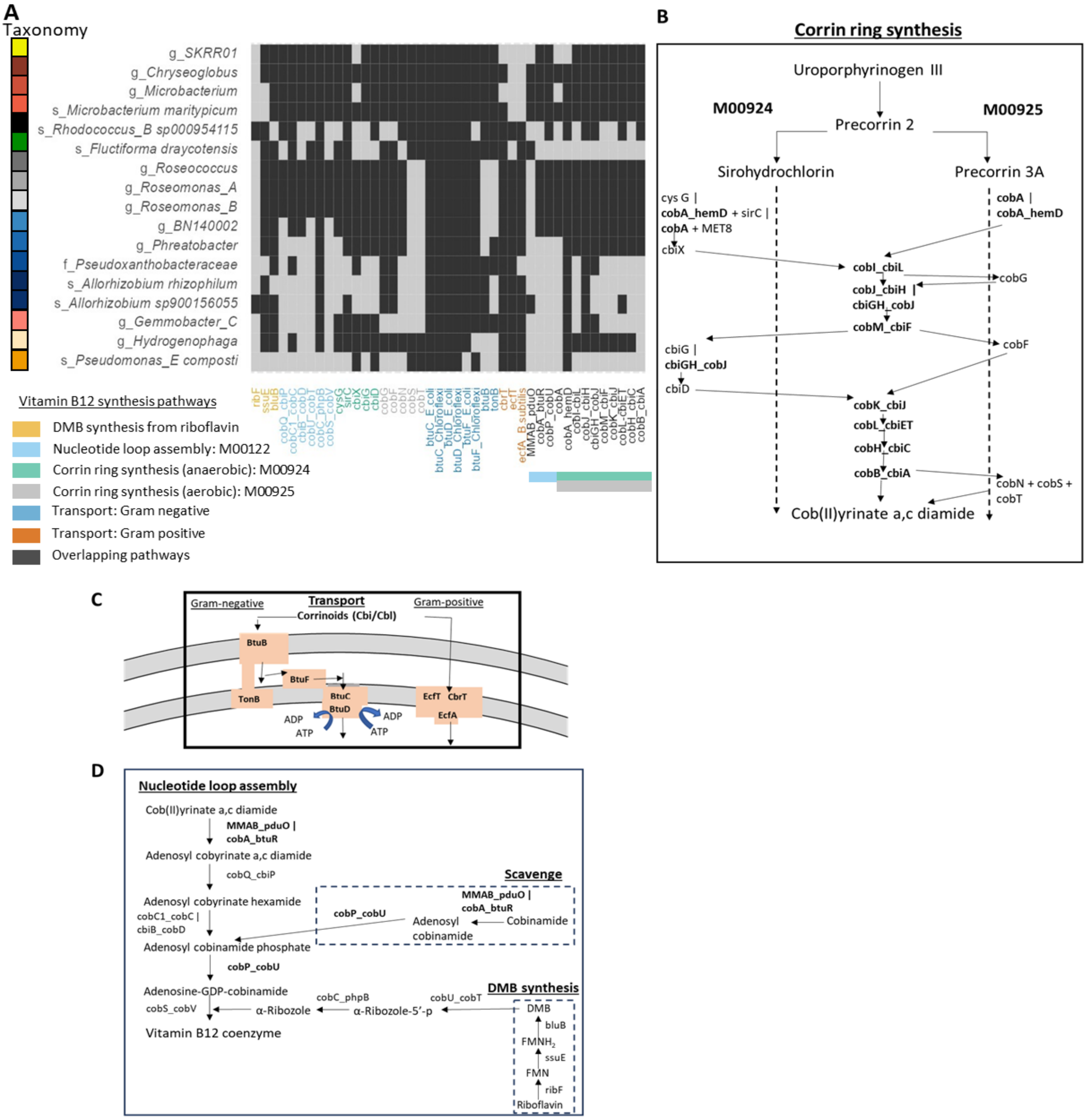


**Figure S9. Analysis of vitamin B12 (cobalamin) biosynthetic and transport pathways.** Representation of completeness of vitamin B12 pathways defined on the KEGG database (24). **(A)** Heatmap plot showing presence (light grey) and absence (dark grey) of select genes, colour-coded by pathway, across the final set of 17 species. See Table 1 for full taxonomy. See *Methods* for full details of gene annotation and analysis and *Supplementary file* 4 for KO analyses. Overlapping pathway genes are those found in more than one pathway, shown by the colour-coded strips. Genes encoding the scavenging pathway directly overlap with genes required for nucleotide loop assembly **(D)**. **(B)** Upper pathway for vitamin B12 synthesis: corrin ring synthesis, defined for both anaerobic and aerobic pathways. Genes involved in these multi-step pathways are listed separately for unique genes. Overlapping genes are highlighted in bold and those used for identical conversions are presented in the centre of the two pathways. Steps in the pathways with alternative gene options are shown with gene names separated by vertical bars. **(C)** Transport pathways of corrinoids (Cbi = cobinamide;

Cbl = cobalamin) across the cell membrane(s) for Gram-negative and Gram-positive bacteria respectively, illustrating those described in (28). **(D)** The lower pathway of vitamin B12 synthesis as nucleotide loop assembly. This pathway is downstream of **(B).**

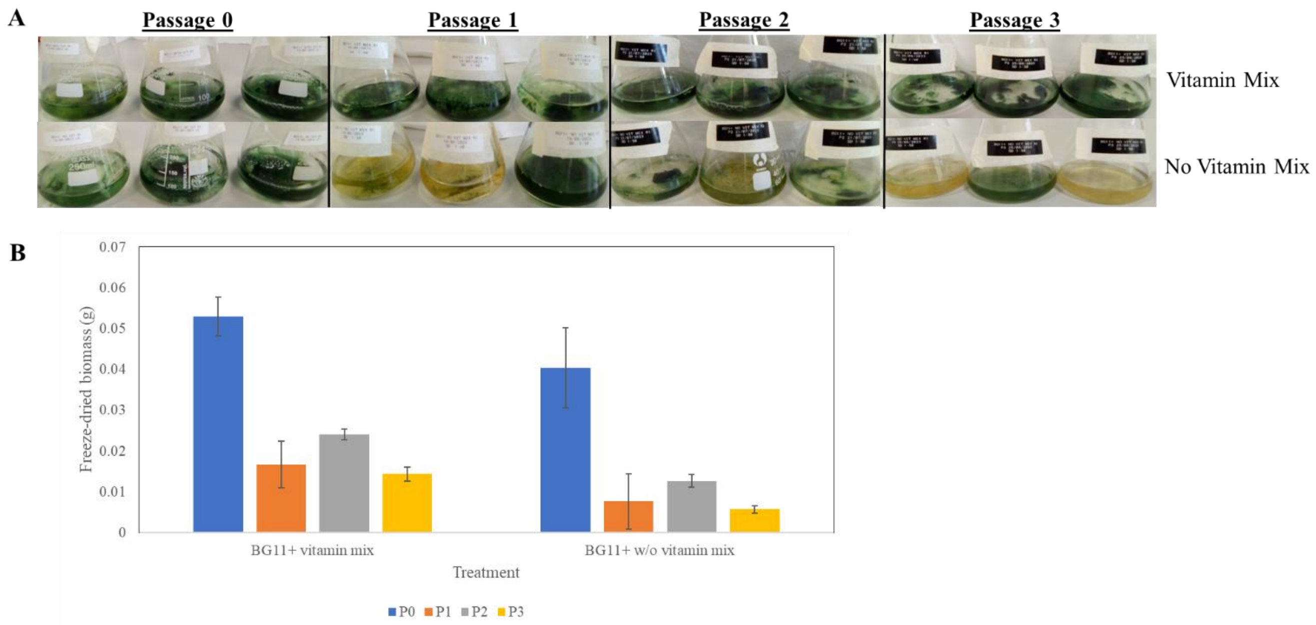


**Figure S10. Growth of community cultures in BG11+ medium with and without vitamin mix supplementation. (A)** Photos showing growth of three replicate cultures after 49 days growth, grown with or without vitamin mix addition, over four serial growth cycles (named Passages 0 - 3). (**B**) Total biomass weight as freeze-dried material per passage measured after 49 days of growth. N = 3 per treatment and passage with Standard Error bars shown. Note that biomass was greater for the first passage for both treatments due to a larger culture volume (see *SI Methods*).

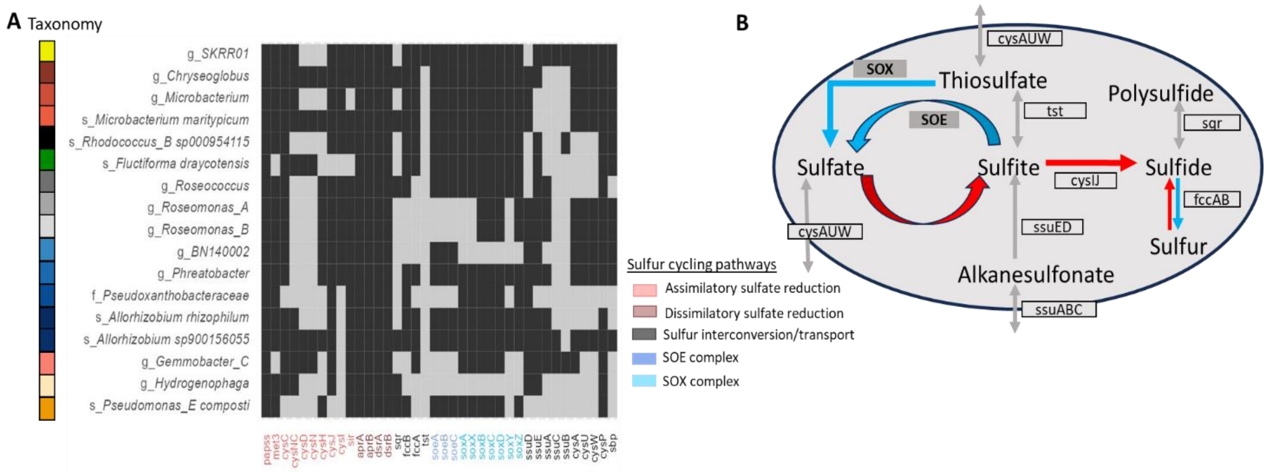


**Figure S11. Functional potential for sulfur metabolism from long and short-read genomes. (A)** Heatmap representation of gene presence (light grey) and absence (dark grey) for major genes encoding sulfur compound assimilation, reduction, interconversion, transport and oxidation pathways, as defined on the KEGG database (24). Full taxonomy is described in Table 1. Bins are colour-categorised as in Fig. 2 and named by the highest taxonomic resolution resolved per bin (genus or species). **(B)** A schematic representation of the key sulfur pathways. The oval outline represents the cell membrane, and uptake or secretion of sulphur compounds as well as their interconversions indicated with grey arrows. Blue and red coloured arrows indicate sulphur-compound oxidation and reduction respectively. Some genes analysed were absent in all genomes, particularly those encoding the dissimilatory sulfate reduction pathway. For a full list of genes (KOs) analysed, see *Supplementary file* 5.

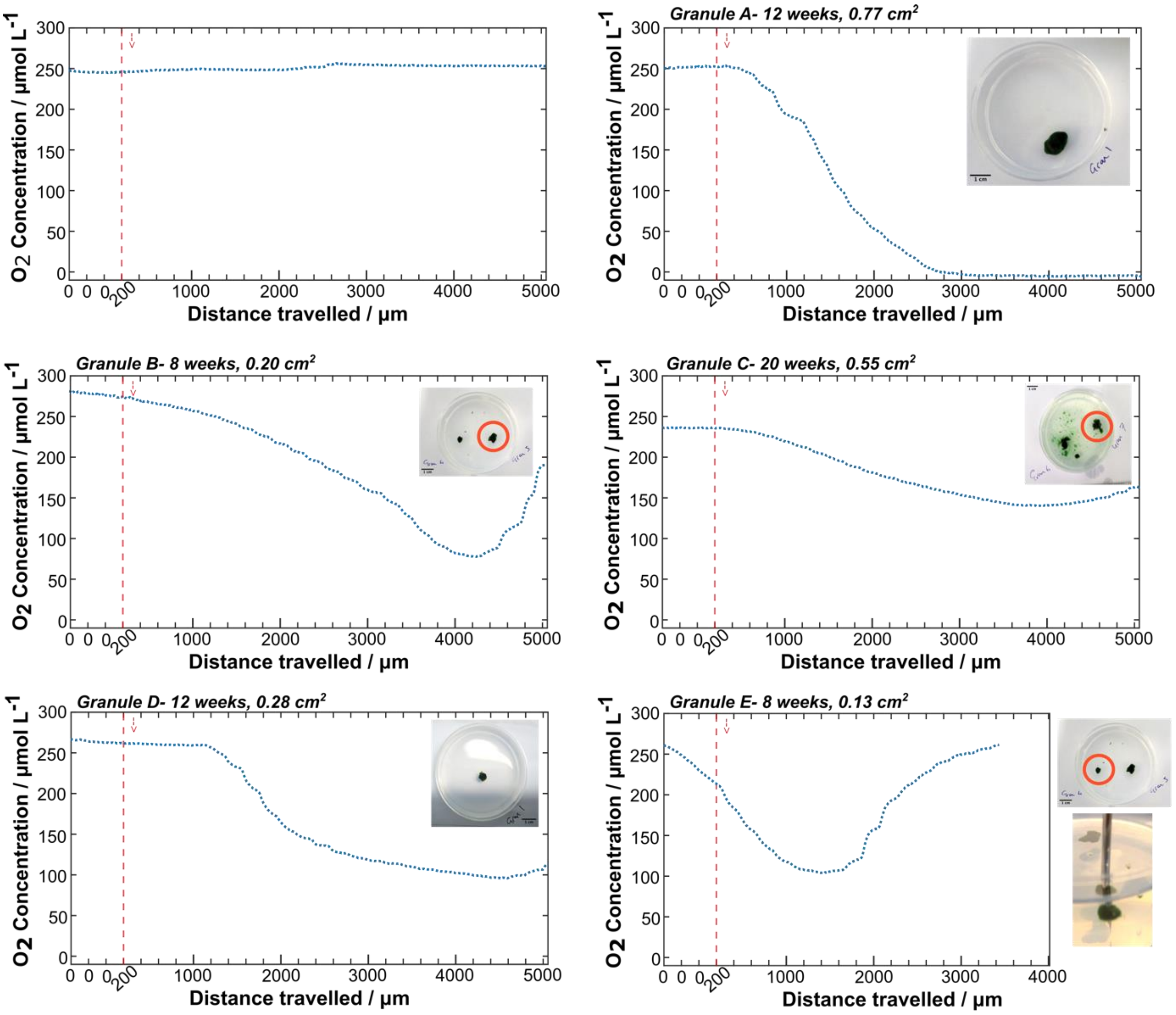


**Figure S12. Additional *Z*-axis oxygen profiles of granules.** Similar profiles to Fig. 3D, across granules of different size and age. 0 µm represents the starting position 1 mm above the granule, followed by the distance travelled through the granule following a timed z-axis step program (200 µm step, at each 20 sec point, unless otherwise stated). Following an initial recorded 60 second rest to allow the signal to stabilise, the red vertical line shows the first descending step of the experiment. The inset images show the investigated granule corresponding with each profile, some circled in red. The top left figure shows the oxygen profile where the experiment procedure is performed descending into the agar bed, instead of into a granule. This steady oxygen concentration measured demonstrates that the oxygen change is caused by the inside granule environment, and not from the probe moving from the BG11+ vitamin mix liquid medium into a semi-solid environment. Granules B and E show how for smaller granules, the probe can pass through the whole granule and into the agar, resulting in the oxygen concentration increasing once it has passed into the agar (as shown in the inset image where the probe has fully passed through the granule). The largest granule, Granule A, is sufficiently large that the core of the granule is fully anoxic, reaching an oxygen concentration of effectively 0 µmol $L^{-1}$. Granules were sampled from the following equivalent cultures: Granule A = P24, Granule B = P25, Granule C = P23, Granule D = P23, Granule E = P25.

### SUPPLEMENTARY TABLES

**Table S1. Comparison of short-read coverage values per species in a live community versus a cryo-revived community sample.** Bin (taxon) coverage values normalised per Gbp of sequence data. P11 = Passage 11, representing a later sub-culture from continued passaging of the community culture (see *Methods*). "P6 cryo" represents the revived cryostock prepared in 10% glycerol of the P6 community presented in Fig 2.

| **Taxon** | **P_11** | **P_6_cryo** |
|---|---|---|
| g_*SKRR01* | 0.078152 | 0.010678 |
| g_*Chryseoglobus* | 0.770111 | 0.951183 |
| g_*Microbacterium* | 0 | 0 |
| s_*Microbacterium maritypicum* | 0.02565 | 0.078846 |
| s_*Rhodococcus_B sp000954115* | 0 | 0 |
| s_*Fluctiforma draycotensis* | 193.3687 | 193.2562 |
| g_*Roseococcus* | 0.134499 | 0.465862 |
| g_*Roseomonas_A* | 0.421759 | 0.444892 |
| g_*Roseomonas_B* | 0.502554 | 0.445647 |
| g_*BN140002* | 6.65E-06 | 6.93E-06 |
| g_*Phreatobacter* | 0.915483 | 1.908237 |
| f_*Pseudoxanthobacteraceae* | 0.578364 | 1.439356 |
| g_*Allorhizobium* | 0.421458 | 0.867304 |
| g_*Gemmobacter_C* | 0.599509 | 0.679752 |
| g_*Hydrogenophaga* | 0.920889 | 2.151485 |
| s_*Pseudomonas_E composti* | 7.136439 | 3.407147 |

**Table S2. Bin normalised coverage depth per giga-base pair (Gbp) and correlations across passages.** Bin coverage values normalised per Gbp of sequencing (as presented in Fig. 2A) were analysed for their correlation with sample time (see *Methods* for full details). See Table 1 for taxonomy of bins. To perform Pearson correlation of each bin/MAG with passage number, a log-10 transformation was applied to the normalised coverages. For Passage 6, no coverage of the *Rhodococcus_B_sp000954115* bin was detected so a pseudo-coverage value of 1.37e-06 (one order of magnitude lower than the smallest recorded coverage value) was applied to that Passage 6 bin. See Fig. 2B for a heatmap presentation of this transformed data. Pearson's correlation r coefficient and p-values are presented, the latter adjusted with the Benjamini-Hochberg correction.

| **Bin** | **P0** | **P3** | **P4** | **P5** | **P6** | **Pearson's Correlation r coefficient** | **Benjamini-Hochberg adjusted p-value** |
|---|---|---|---|---|---|---|---|
| g_*SKRR01* | 0.531 | 0.409 | 0.704 | 0.980 | 0.401 | 0.21 | 0.847 |
| g_*Chryseoglobus* | 1.670 | 1.314 | 1.675 | 1.451 | 1.393 | -0.5 | 0.626 |
| g_*Microbacterium* | -1.012 | -1.184 | -3.253 | -4.866 | -4.468 | -0.87 | 0.28 |
| s_*Microbacterium maritypicum* | 0.358 | -0.287 | 0.356 | 0.490 | 0.826 | 0.46 | 0.641 |
| s_*Rhodococcus_B sp000954115* | -0.305 | -1.492 | -1.506 | -0.858 | -5.866 | -0.68 | 0.366 |
| s_*Fluctiforma draycotensis* | 2.052 | 2.108 | 2.021 | 1.836 | 1.879 | -0.71 | 0.361 |
| g_*Roseococcus* | -0.295 | 0.260 | -0.00759 | -0.238 | -0.379 | -0.14 | 0.874 |
| g_*Roseomonas_A* | 0.723 | 0.278 | 0.00725 | 0.755 | 0.624 | -0.07 | 0.91 |
| g_*Roseomonas_B* | -0.470 | -0.715 | -0.304 | 0.467 | 0.237 | 0.71 | 0.361 |
| g_*BN140002* | 0.499 | 1.195 | -4.609 | -4.338 | -4.540 | -0.79 | 0.361 |
| g_*Phreatobacter* | 1.231 | -0.0302 | 0.843 | 1.355 | 1.393 | 0.21 | 0.847 |
| f_*Pseudoxanthobacteraceae* | 0.541 | 0.538 | 0.681 | 1.343 | 1.0792 | 0.75 | 0.361 |
| g_*Allorhizobium* | 0.446 | 0.708 | 0.888 | 1.0426 | 0.870 | 0.9 | 0.28 |
| g_*Gemmobacter_C* | 0.590 | 0.820 | 0.608 | 0.795 | 0.918 | 0.72 | 0.361 |
| g_*Hydrogenophaga* | 0.334 | 0.717 | 0.880 | 0.712 | 1.101 | 0.91 | 0.28 |
| s_*Pseudomonas_E composti* | 1.0394 | 0.815 | 1.221 | 0.956 | 1.179 | 0.28 | 0.847 |

**Table S3. Assembly and genome statistics for the 17-species bacterial community.** Statistics were collected from annotation of MAGs with the DFAST pipeline (19). PacBio long-read MAGs were analysed for all species, apart from MAGs 3 and 5 (labelled with *) for which only short-read MAGs were available. MAG completeness score represents the percentage of the total of 36 single copy core genes (SCGs) that are present, based on SCG clustering at 99% sequence similarity level (see *Methods*). Completeness was calculated using CheckM for MAG numbers 2, 3, 5 and 6.

| MAG | Taxon | No. contigs | N50 (bp) | Genome size (bp) | GC content (%) | No. coding seqs | Completeness |
|---|---|---|---|---|---|---|---|
| 1 | g_*SKRR01* | 8 | 4240744 | 4350865 | 71.3 | 4030 | 100 |
| 2 | g_*Chryseoglobus* | 1 | 2902611 | 2902611 | 65.8 | 2720 | 98.4 |
| 3 * | g_*Microbacterium* | 21 | 279023 | 3753668 | 68.5 | 3489 | 98.04 |
| 4 | s_*Microbacterium maritypicum* | 1 | 3640567 | 3640567 | 68.4 | 3447 | 100 |
| 5 * | s_*Rhodococcus_B sp000954115* | 1533 | 4110 | 5152295 | 64.9 | 3638 | 88.23 |

| | | | | | | | |
|---|---|---|---|---|---|---|---|
| 6 | s_*Fluctiforma draycotensis gen. nov., sp. nov.* | 1 | 4790552 | 4790552 | 44.3 | 4360 | 98 |
| 7 | g_*Roseococcus* | 9 | 4547859 | 4705185 | 71.2 | 4386 | 100 |
| 8 | g_*Roseomonas_A* | 1 | 6768607 | 6768607 | 70.8 | 6344 | 100 |
| 9 | g_*Roseomonas_B* | 16 | 3660211 | 5674990 | 71.3 | 5364 | 100 |
| 10 | g_*BN140002* | 22 | 412864 | 5357319 | 67.3 | 5265 | 75 |
| 11 | g_*Phreatobacter* | 13 | 1991051 | 4478198 | 68.8 | 4235 | 100 |
| 12 | f_*Pseudoxanthobacteraceae* | 7 | 4010775 | 5524412 | 70.3 | 5021 | 100 |
| 13_strain 1 | s_*Allorhizobium rhizophilum* | 75 | 308893 | 3807890 | 61.7 | 3685 | 80.6 |
| 13_strain 2 | s_*Allorhizobium rhizophilum* | 138 | 315789 | 6718987 | 61.6 | 6322 | 100 |
| 14 | s_*Allorhizobium sp900156055* | 92 | 74608 | 4716025 | 61.3 | 5343 | 88.9 |
| 15 | g_*Gemmobacter_C* | 41 | 4375673 | 5359230 | 66.6 | 4937 | 100 |
| 16 | g_*Hydrogenophaga* | 57 | 74981 | 3754005 | 65.6 | 4002 | 91.7 |
| 17 | s_*Pseudomonas_E composti* | 1 | 5235516 | 5235516 | 62.5 | 4793 | 100 |

**Table S4. Salt solution composition of minimal medium (MM)**. This recipe is based on (1) and has been altered to include vitamin and trace metal mixtures (described below).

| salt name | g/mol | g/L | mM |
|---|---|---|---|
| Potassium nitrate | 101.10 | 5 | 49.4544 |
| Dipotassium phosphate | 174.18 | 0.1145 | 0.6573 |
| Magnesium sulfate | 120.37 | 0.02 | 0.1662 |
| Iron (III) chloride hexahydrate | 270.30 | 0.00833 | 0.0308 |
| EDTA disodium salt dihydrate | 372.24 | 0.1107 | 0.2974 |

**Table S5. Trace metals composition for MM.** This trace metal solution is prepared as a hundred times concentrated stock solution.

| Trace Metal | g/L(stock) | g/L medium | g/mol | mol/l (medium) |
|---|---|---|---|---|
| Nitrilotriacetic acid | 1.5000 | 0.015000 | 191.14 | 7.8E-05 |
| Magnesium chloride hexahydrate | 2.4800 | 0.024800 | 203.3 | 1.2E-04 |
| Manganese chloride tetrahydrate | 0.5854 | 0.005854 | 197.9 | 3.0E-05 |
| Sodium chloride | 1.0000 | 0.010000 | 58.44 | 1.7E-04 |
| Iron chloride tetrahydrate | 0.0715 | 0.000715 | 198.81 | 3.6E-06 |
| Cobalt chloride hexahydrate | 0.1524 | 0.001524 | 237.93 | 6.4E-06 |
| Calcium chloride dihydrate | 0.1000 | 0.001000 | 147.01 | 6.8E-06 |
| Zinc chloride tetrahydrate | 0.0853 | 0.000853 | 136.315 | 6.3E-06 |

| | | | | |
|---|---|---|---|---|
| Copper chloride | 0.0054 | 0.000054 | 134.452 | 4.0E-07 |
| Aluminium chloride | 0.0103 | 0.000103 | 133.34 | 7.7E-07 |
| Boric acid | 0.0100 | 0.000100 | 61.83 | 1.6E-06 |
| Disodium molybdate dihydrate | 0.0100 | 0.000100 | 205.92 | 4.9E-07 |
| Nickel chloride hexahydrate | 0.0300 | 0.000300 | 237.69 | 1.3E-06 |
| Disodium selenite pentahydrate | 0.0003 | 0.000003 | 262.94 | 1.1E-08 |
| Sodium tungstate dihydrate | 0.0080 | 0.000080 | 329.8477 | 2.4E-07 |

**Table S6. Vitamin mix composition for MM and for addition to BG11+ medium.** This vitamin mix solution is prepared as a thousand times concentrated stock solution.

| **Vitamins** | **g/L(stock)** | **g/L medium** | **g/mol** | **mol/l (medium)** |
|---|---|---|---|---|
| Biotin | 0.020 | 0.00002 | 244.31 | 8.2E-08 |
| Folic acid | 0.020 | 0.00002 | 441.4 | 4.5E-08 |
| Pyridoxine hydrochloride | 0.100 | 0.0001 | 205.63 | 4.9E-07 |
| Thiamine hydrochloride | 0.050 | 0.00005 | 337.26 | 1.5E-07 |
| Riboflavin | 0.050 | 0.00005 | 376.36 | 1.3E-07 |
| Nicotinic acid | 0.050 | 0.00005 | 123.11 | 4.1E-07 |
| D-Calcium pantothenate | 0.050 | 0.00005 | 238.265 | 2.1E-07 |
| para-Aminobenzoic acid | 0.050 | 0.00005 | 137.14 | 3.6E-07 |
| Cobalamin | 0.001 | 0.000001 | 1355.37 | 7.4E-10 |
| Lipoic acid | 0.050 | 0.00005 | 206.33 | 2.4E-07 |
